# Twin Study of Early-Onset Major Depression Finds DNA Methylation Enrichment for Neurodevelopmental Genes

**DOI:** 10.1101/422345

**Authors:** Roxann Roberson-Nay, Aaron R. Wolen, Dana M. Lapato, Eva E. Lancaster, Bradley T. Webb, Bradley Verhulst, John M. Hettema, Timothy P. York

**Author notes:** Correspondence: Roxann Roberson-Nay, Ph.D., Virginia Commonwealth University, Department of Psychiatry, Virginia Institute for Psychiatric and Behavioral Genetics, P.O. Box 980489, Richmond, VA 23298.

## Abstract

Major depression (MD) is a debilitating mental health condition with peak prevalence occurring early in life. Genome-wide examination of DNA methylation (DNAm) offers an attractive complement to studies of allelic risk given it can reflect the combined influence of genes and environment. The current study used a co-twin control design to identify differentially and variably methylated regions of the genome that distinguish monozygotic (MZ) twins with and without a lifetime history of early-onset MD. The sample included 150 Caucasian monozygotic twins (73% female; *M*age=17.52 *SD*=1.28) assessed during a developmental stage characterized by relatively distinct neurophysiological changes. All twins were generally healthy and currently free of medications with psychotropic effects. DNAm was measured in peripheral blood cells using the Infinium Human BeadChip 450K Array. MD associations were detected at 760 differentially and variably methylated probes/regions that mapped to 428 genes. Results indicated an association between early-onset MD and many genes and genomic regions involved in neural circuitry formation, projection, functioning, and plasticity. Gene enrichment analyses implicated genes related to neuron structures and neurodevelopmental processes including cell-cell adhesion genes (e.g., *CDHs, PCDHAs, PCDHA1C/2C*). Genes previously implicated in mood and psychiatric disorders as well as chronic stress (e.g., *HDAC4, NRG1*) also were identified. DNAm regions associated with MD where found to overlap genetic loci observed in the latest Psychiatric Genomics Consortium meta‐analysis of depression. Understanding the time course of epigenetic influences during emerging adulthood may clarify developmental phases where genes modulate individual differences in MD risk.

## Introduction

Major depression (MD) is highly prevalent, ranking second in the global burden of disease, with the overall lifetime risk estimated to be 16.2% in the general population^1^. MD is associated with increased mortality, particularly suicide^1^. Among adolescents, MD is associated with the greatest level of impairment of all psychiatric conditions, with 16% of females and 12% of males endorsing at least one major depressive episode (MDE) by age 18^2^. An early age of onset confers increased risk for negative socioemotional outcomes including recurrent MDEs^3,4^. Adolescence/young adulthood is characterized by neurophysiological changes (e.g., synaptic pruning, myelination) that significantly influence brain function and behavior, which may increase risk for MD and other psychiatric conditions^5^. Thus, understanding the genetic contributions to MD during this dynamic neurophysiological period where peak incidence is observed^2,6–8^ is critical to elucidating developmentally informed pathways to mood disorders.

Twin and family studies robustly demonstrate that genetic factors play a role in risk for MD, with heritability estimates of roughly 35% for MD and 45% for early-onset MD^9^. The largest meta-analysis of molecular genetic studies of MD recently identified 44 independent loci, underscoring the centrality of genetic factors in the etiology of MD^10^. Twin study variance component analyses also indicate a considerable contribution of unique environmental risk factors to MD^11^. Due to the substantial link between environmental hardship and onset of a MDE^12,13^, epigenetic mechanisms may, in part, mediate the influence of environmental stress and interact with genetic liability for MD over the lifespan^14–16^ that can influence the expression of genes but do not alter the underlying genetic sequence. Animal studies have been critical to demonstrating a causal association between early life environments, epigenetic alterations, and phenotypic outcomes. For example, the seminal work of Michael Meaney and his research team demonstrates the importance of maternal care in altering the expression of genes that regulate behavioral and neuroendocrine responses to stress as well as synaptic development in the rat hippocampus^17–22^. Indeed, a number of animal and ^16^. Epigenetic mechanisms refer to DNA, chromatin, and RNA human studies demonstrate lasting epigenetic alterations occurring in the genomes of cells including changes to post-mitotic neurons that integrate experience-dependent changes^23^. Thus, the timing of environmental stress plays an important role in subsequent epigenetic consequences, with early life stress paradigms in mice and humans demonstrating enduring changes in epigenetic profiles^17,20,24–28^.

A number of studies have utilized genome-wide platforms to determine DNA methylation (DNAm) differences between MD cases and controls. However, as much as 37% of methylation variance can be accounted for by genetic factors^30^ with recent studies indicating that common genetic variation (i.e., methylation quantitative trait loci [*<u>mQTLs</u>*]) influence DNAm levels^31–35^. Most MD case-control studies of DNAm do not account for allelic variation which means genetic and environmental influences on DNAm cannot be disaggreagated. In contrast, the quasi-experimental design afforded by the co-twin control approach greatly improves on the unmatched case‐control design (see Supplemental Figure 1). The use of monozygotic (MZ) twins removes the impact of unmeasured confounds such as genetic variation, uterine environment, age, sex, race, cohort effects, and exposure to many shared environmental events.

The current study utilitized the robust co-twin control method and a statistically powerful approach to detect differentially and variably methylated DNAm regions associated with MD in a sample of adolescent and emerging adult twins. Studying this developmental period offers a number of advantages over later life periods including fewer confounds to DNAm variability such as a history of prolonged or multiple psychiatric/medical comorbidities and medication usage as well as long-term nicotine use. Moreover, it eliminates the well-known epigenetic changes associated with aging^36–38^. The developmental window of young adulthood also is associated with moderate conservation of DNAm that is nonetheless responsive to environmental signals^39^, making it an ideal sensitive period for the study of MD.

## Methods

### Participants

One hundred sixty-six MZ twins (83 pairs) were selected from a larger sample of twins (N=430 pairs) enrolled in a study examining risk factors for internalizing disorders in a general population sample of twins (R01MH101518)^40^. Twins were primarily recruited through the Mid‐Atlantic Twin Registry (MATR), a population-based registry^40,41^. All participants (parents/minor children, adults) provided written informed consent/assent.

Of the 83 twin pairs initially identified as MZ via the zygosity questionnaire, five pairs (6.3%) were determined to be DZ pairs using DNA-based markers and were removed (for zygosity determination, see Supplemental). One or both twins from three additional pairs failed DNA-based quality control checks, reducing the final analyzed sample to 75 MZ twin pairs (150 twins; see Table 1). All twins were raised together in the same home and were required to be free of psychotropic medications/medications with psychotropic effects at the time of study entry, although approximately 4.0% (n=6) endorsed a history of psychotropic medication use, slightly lower than the national average^42^. See full study exclusionary criteria in Supplement.

**Table 1.** Demographic and clinical characteristics of twins meeting definite or probable DSM-5 criteria for lifetime history of MD (MD Affected) versus no lifetime history of MD (MD Unaffected).

| | *M* (SD) or n (%) | | *t*/$\chi^2$ | *p* |
| --- | --- | --- | --- | --- |
|  | MD Unaffected<br>n=111 | MD Affected<br>n=39 |  |  |
| **Demographic/Sample** |  |  |  |  |
| Age, years | 17.49 (1.3) | 17.60 (1.3) | 0.66 | 0.51 |
| Sex, Female | 81 (73.0%) | 29 (74.4%) | 0.28 | 0.87 |
| Ethnicity, Hispanic | 5 (4.5%) | 3 (7.7%) | 0.58 | 0.43 |
| Nicotine Use<sup>§</sup>, Current Smoker | 2 (1.8%) | 3 (7.7%) | 3.11 | 0.11 |
| **Clinical Characteristics** |  |  |  |  |
| SMFQ | 4.4 (3.7) | 9.0 (6.0) | 6.40 | <.001 |
| History of psychotropic medication use | 2 (1.8%) | 4 (10.3%) | 0.64 | 0.62 |
| Panic Disorder | 5 (4.5%) | 3 (7.7%) | 0.56 | 0.43 |
| Social Anxiety Disorder | 10 (9.1%) | 8 (20.5%) | 3.54 | 0.08 |
| Specific Phobia | 8 (7.3%) | 7 (17.9%) | 3.63 | 0.07 |
| Generalized Anxiety Disorder | 2 (1.8%) | 4 (10.3%) | 5.31 | 0.04 |
| **MD Features** |  |  |  |  |
| Age of Onset (years) | --- | 14.9 (1.7) |  |  |
| Number of Major Depressive Episodes, n |  |  |  |  |
| 1 episode | --- | 17 (43.6%) |  |  |
| 2-3 episodes | --- | 16 (41.0%) |  |  |
| 3-5 episodes | --- | 3 (7.7%) |  |  |
| ≥6 episodes | --- | 3 (7.7%) |  |  |
| Number of symptoms during worst MDE | --- | 5.69 (1.1) |  |  |

*Note:* SMFQ=Short Mood and Feelings Questionnaire.<sup>§</sup>Fagerstrom Test for Nicotine Dependence score ranged 2 (low dependence) to 6 (moderate dependence) with Mode=2, Median=2, and Mean=3.

### Measures

#### Major Depression

All twin pairs completed a psychiatric history based on an expanded version of the Composite International Diagnostic Interview (CIDI)-Short Form, which queried DSM-5 MD Criterion A and C (see Supplement for questions and diagnostic algorithm)^43^. Current depressive symptoms also were assessed using the Short Mood and Feelings Questionniare (SMFQ), which is a 13-item questionnaire validated to measure depressive symptoms in adolescents and adults^44^.

### DNA Methylation Processing

Detailed information concerning DNA extraction, methylation assessment, normalization, and quality control (QC) can be found in the Supplement.

### DNA Methylation Analysis

#### Analytic Approach

Based on observations that the average correlation between probes on the 450K microarray within approximately 250 base pairs (bp) is 0.83 and within 1 kb is 0.45^45–47^, several methods have been proposed to take advantage of this structure to identify consistent DNAm change across a contiguous region^45,46,48,49^. Current approaches quantify regional DNAm change either as a mean difference (differentially methylated region [DMR]) or as a difference in variance (variably methylated region [VMR]). Due to the sparse and highly clustered placement of features on the Illumina 450K platform, a custom approach for defining DMRs and VMRs was developed for this study motivated by the algorithm proposed by Ong et al. ^45,50^. They suggest both a regional and individual CpG probe approach run in tandem since approximately 25% of probes do not have a neighboring probe within 1 kb. To this end, the single-probe analysis provided the raw materials for the regional approach adopted to identify and assess significance of DMRs and VMRs. Type I error rates were estimated from empirical *P*-values for all univariate statisitics (mean level and variance based tests) described below were calculated using a permutation approach. For *k* = 1,000 reorderings, the outcome variable was resampled in a way that preserved the discordance/concordance pair status frequencies. The false discovery rate (FDR)^51^ was estimated from the distribution of these empirical *P*-values.

### Identifying Regional DNAm Change

#### Differentially Methylated Regions

Potential DMRs were constructed using only those individual CpG probes associated with MD status resulting in a test statistic in the upper or lower 5^th^ percentile of all probes tested. Univariate tests were performed by fitting a linear mixed-effects model^52^ separately for each probe by regressing the normalized DNAm probe intensity on MD status while adjusting NK cell proportion and inclusion of a random effect term to account for twin pair membership. NK cell proportion was the only estimated cell type nominally significantly different between MD cases and controls (*t* = 0.266, *p* = 0.051), so it was included as a covariate to control for potential bias that might arise from DNAm differences due to changes in NK blood cell proportions rather than those attributable to DNAm changes associated with MD itself.

The structure of MZ twin exposure groupings provided an opportunity to focus on DNAm associations that are unique to MD versus other experimental design features. The sample used for study was composed of three types of MZ twin pairs (i.e., discordant MD, concordant MD, concordant healthy) and resulted in number of potential statistical contrasts to be tested (Supplemental Table 1). Specific contrasts were simultaneously fit within the linear model to estimate the degree of association among two sets of comparisons: 1) the effect of interest (e.g., MD affected versus MD unaffected) and 2) extraneous associations not of interest. The latter set of contrasts, while of general interest, do not directly relate to MD and were filtered from downstream applications. For example, the comparison of DNAm levels where MD is expressed in both members (concordant MD) versus only a single member (discordant MD) may be associated with a unique DNAm profile (i.e., line 2 in Supplemental Table 1). Probes with test statistics in the 1^st^ and 99^th^ quantiles identified in any contrast not of interest were removed.

From the set of filtered CpG probes, a candidate DMR was defined as having at least 2 contiguous CpGs within 1 kb. The strength of a DMR association was estimated by a test statistic reflecting the area of influence approximated by the trapezoidal rule where the height (*h*) was the length in base pairs between two contiguous CpGs and *a* and *b* were the univariate test statistic for the contiguous CpGs. For DMRs with more than two CpGs, the area for each contiguous pairing was summed to represent the area under the curve (AUC) for the entire DMR. All probes in the DMR had the restriction of test statistics with the same sign. A positive test statistic indicated hypermethylation in cases versus controls while a negative test statistic indicated hypomethylation.

#### Variably Methylated Regions

A similar strategy was adopted to identify variably methylated regions (VMR). In this case, the test statistic calculated was the *F*-value comparing the variance of two samples applied to the filtered set of CpG probes. Again, only probes for the contrast of interest were considered (10^th^ / 90^th^ percentiles) if they did not intersect with those from contrasts not of interest (1^st^/99^th^ percentiles). VMRs were restricted to at least two contiguous probes within 1 kb whereby all probes had the restriction of either increasing or decreasing variability in cases versus controls.

#### Significance Assessment of Regional Change

The statistical significance of DMR and VMR AUC estimates were assessed using a rank-based permutation method^53^. This nonparametric method estimates an FDR without relying on strong assumptions about the normality of the data. The *k* = 1,000 permutations of univariate tests were used to estimate the expected order statistics. The FDR was calculated based on the observed versus expected null scores. Briefly, for a range of thresholds, regions are called significant if the value of the observed ordered test statistic minus the mean value from the permuted rank exceeds a given threshold. The number of falsely called regions is the median number of regions that exceed the lowest AUC value of regions called significant. The FDR is calculated as the ratio of the number of falsely called regions to the number of regions called significant. An implementation of the SAM algorithm is available as a R package^54^ but was recoded to allow for flexibility in specifying models for the twin data and to trim extreme test statistics likely to be false positives before the calculation of the FDR.

### Functional and Regulatory Enrichment

The distribution of significant CpG probes and regions identified to be differentially and variably methylated by MD status were examined separately across functional and regulatory annotations. CpG findings were mapped to known genes^55^ for enrichment of Gene Ontology classifications^56^ using clusterProfiler^57^. Classification functons included biological processes, cellular components, and molecular function, in addition to KEGG pathways. Tests for non random association of CpG island features and ChromHMM chromatin states were based on the AH5086 and AH46969 tracks from the AnnotationHub package^58^, respectively. CpG island shores were defined as being 2 kb regions flanking CpG islands while shelves were demarcated as 2 kb upstream or downsteam shore regions. A test of enrichment for each of these annotations was calculated by comparing the proportion of sequence from the intersection of significant CpG regions with the regions defined by the annotation feature. Bootstrap methods using 1,000 resamplings were used to estimate 95% confidence intervals. This observed overlap was compared to an empirical distribution of random samples of genome groups of the same size and structure drawn from the background set under consideration. Empirical P-values were calculated from 1,000 random reorderings of the data using standard methods^59^.

### PGC GWAS Enrichment

A similar resampling method was performed to count the number of significant CpG regions that overlapped with the findings of a recent genome-wide association study (GWAS) meta-analysis conducted by the Psychiatric Genomics Consortium (PGC) group^10^. The depression phenotype in this meta-analysis was derived from a number of different methods including clinical interview, self-report, electronic medical record abstraction, and self-report of a lifetime diagnosis. This study identified 44 MD-associated loci across 18 chromosomes, which included genes enriched for targets of antidepressant medication. The non-random frequency of overlap between the significant CpG regions and the 44 independent PGC findings was assessed using bootstrap and permutation approaches from 1,000 data resamplings.

## Results

### Sample Characteristics

MZ twins meeting DSM-5 MD criteria at the probable or definite level self-reported higher depression symptom scores on the SMFQ and had higher rates of generalized anxiety disorder compared to MD unaffected twins (*t*(1,146)=6.4, *p*<.001); *χ*^2^(1)=5.31, *p*=.04, respectively; see Table 1). For those twins meeting DSM-5 criteria for at least one MDE, the mean age at onset was approximately 15 years, and the majority of twins (∼85%) reported experiencing 3 or fewer MDEs in their lifetime. Most MD affected twins (82%) reported five or more symptoms during their worst lifetime MDE. Subject age, sex, self-reported ethnicity, and combustible cigarette use did not differ by MD status.

### Differentially Methylated Probes and Regions

From the set of 455,828 screened CpG probes, according to our definition, 50,990 background regions could be created, covering 59.9 megabases. After DMP tests, 3,995 regions consisting of 28,600 CpG probes could be considered candidate DMRs. Seventeen DMRs were identified as significantly associated with MD (all hypermethylated in MD cases) of which 15 mapped onto genes (FDR 9.9%; see Table 2). The number of CpG probes in significant DMRs ranged from 3 to 7 (median=4). Individual probe testing (DMP) resulted in 59 hypomethylated and 77 hypermethylated CpG sites with respect to MD status (FDR 1%; Supplemental Table 2). The combined set of 30.6 kb DNA sequence covered by significant DMR and DMP findings was found to have a nonrandom pattern of enrichment across ChromHMM annotations, specifically sites of strong transcription (*p*=0.019), enhancers (*p*=0.045), ZNF genes/repeats (*p*=0.001), heterochromatin (*p*=0.013) and weak repressed polycomb (*p*=0.045) (Supplemental Figure 2) and CpG island relationships which included both north (*p*=0.024) and south (*p*=0.003) shelf regions (Supplemental Figure 3).

**Table 2.** Differentially methylated regions (DMRs) where MD affected twins exhibited higher means compared to MD unaffected twins.

| **Chromosome** | **Start** | **End** | **Symbol** | **EntrezID** | **Number_CpG** | **Empirical_AUC** |
| --- | --- | --- | --- | --- | --- | --- |
| chr1 | 36787678 | 36789401 | SH3D21 | 79729 | 3 | 3836.3652 |
| chr1 | 221053841 | 221055665 | HLX | 3142 | 4 | 4242.3503 |
| chr2 | 240035107 | 240036791 | HDAC4 | 9759 | 3 | 4057.6444 |
| chr5 | 1090741 | 1092417 | SLC12A7 | 10723 | 4 | 3818.3311 |
| chr5 | 171709917 | 171711524 | UBTD2 | 92181 | 3 | 4070.7302 |
| chr5 | 176936563 | 176938522 | DOK3 | 79930 | 6 | 3826.3018 |
| chr6 | 30519905 | 30521619 | NA | NA | 5 | 3861.8967 |
| chr6 | 31828260 | 31830030 | NA | NA | 4 | 4558.5396 |
| chr6 | 32797253 | 32798887 | NA | NA | 4 | 3849.6722 |
| chr6 | 39281541 | 39283313 | KCNK17 | 89822 | 3 | 4122.0762 |
| chr6 | 146863647 | 146865487 | RAB32 | 10981 | 7 | 4701.7437 |
| chr16 | 2023998 | 2025868 | TBL3 | 10607 | 4 | 4329.8853 |
| chr16 | 58767249 | 58769104 | GOT2 | 2806 | 4 | 4046.9261 |
| chr17 | 46659019 | 46660940 | HOXB3 | 3213 | 4 | 4501.7814 |
| chr17 | 70116185 | 70118162 | SOX9 | 6662 | 4 | 4120.2233 |
| chr19 | 13213428 | 13215387 | LYL1 | 4066 | 3 | 4381.7764 |
| chr21 | 38069321 | 38070994 | NA | NA | 4 | 3892.4767 |

### Variably Methylated Probes and Regions

Regional analysis identified 10 variably methylated regions (VMRs) signficant from a total of 11,055 candidate VMRs (FDR 17.3%). Seven of the VMRs mapped onto genes (Table 3). Significant VMRs were all more variable in MD cases, and the number of CpG probes in these regions ranged from 2 to 11 (median=4). The VMP analysis yielded 560 significant VMP findings (FDR 1%), all of which were more variable in MD cases except for a single probe (Supplemental Table 3). The combined set of VMR and VMP DNA regions of 16.6 Kb reflected a nonrandom enrichment with ChromHMM annotations for 5’/3’ transcription (*p*= 0.035), genic enhancers (*p*=0.024) and heterochromatin (*p*=0.050) (Supplemental Figure 4) and CpG island relationships within the south shelf (*p*=0.029) (Supplemental Figure 5).

**Table 3.** Variably methylated regions (VMRs) where MD affected twins exhibited greater variance compared to MD unaffected twins.

| **Chromosome** | **Start** | **End** | **Symbol** | **EntrezID** | **Number_CpG** | **Empirical_AUC** |
| --- | --- | --- | --- | --- | --- | --- |
| chr3 | 39321449 | 39323539 | CX3CR1 | 1524 | 4 | 5572.9239 |
| chr3 | 46925081 | 46925524 | PTH1R | 5745 | 2 | 5031.1289 |
| chr3 | 170136920 | 170137321 | CLDN11 | 5010 | 2 | 5110.2437 |
| chr6 | 32049516 | 32049825 | NA | NA | 3 | 5038.8717 |
| chr6 | 169284344 | 169287304 | NA | NA | 5 | 5276.0094 |
| chr6 | 170595385 | 170597898 | DLL1 | 28514 | 8 | 5628.6246 |
| chr7 | 157225062 | 157225567 | NA | NA | 2 | 5131.4817 |
| chr10 | 134739746 | 134741032 | CFAP46 | 54777 | 4 | 5667.4646 |
| chr11 | 64510112 | 64513156 | RASGRP2 | 10235 | 7 | 5458.4387 |
| chr17 | 40837037 | 40839469 | CNTNAP1 | 8506 | 4 | 5035.574 |

### Gene Enrichment Analysis

Genes that mapped to signficant differentially or variably methylated findings were combined for gene-based enrichment to provide an overview of all DNAm contributions at a functional level. The results of enrichment tests yielded significant over-representation for biological processes (BP) and cellular function (CF), and no enrichment for molecular function or KEGG, at the 10% FDR (Table 4). The BP gene category associations were hemophilic cell adhesion and cell‐cell adhesion while the significant terms for CF were associated with functions of neurons, including neuron projection terminus, terminal button, axon part, cell projection part, axon, and presyn apse.

**Table 4.** Gene enrichment analysis summary.

|  | **Ontology Category** | **Description** | **Gene Ratio** | **q-value** | **Gene Symbol** |
| --- | --- | --- | --- | --- | --- |
| GO:0007156 | Biological Process | homophilic cell adhesion via plasma membrane adhesion molecule | 23, 568 | 0.00002 | CDH3, CDH6, PCDHαC1-2, PCDHα1-13, PCDH10, PCDH20, PTPRT, CADM1, PLXNB3, CLSTN2 |
| GO:0098742 | Biological Process | cell-cell adhesion via plasma-membrane adhesion molecules | 28, 568 | 0.00002 | CDH3, CDH6, PTPRT, CLDN4, CADM1, GRID2, ITGA5, CLDN11, PLXNB3, PCDHαC1-2, PCDHα1-13, PCDH20, PTPRD, CLSTN2 |
| GO:0044306 | Cellular Function | neuron projection terminus | 14, 605 | 0.05888 | BAIAP2, AAK1, CYFIP1, SYT11, PACSIN1, ILK, PFN2, PNOC, PVALB, DNAJC5, MOB2, NAPA, BSN, AP3D1 |
| GO:0043195 | Cellular Function | terminal bouton | 9, 605 | 0.05888 | AAK1, CYFIP1, SYT11, ILK, PFN2, PVALB, DNAJC5, NAPA, AP3D1 |
| GO:0033267 | Cellular Function | axon part | 16, 605 | 0.07314 | AAK1, TRAK1, CYFIP1, SYT11, KIF13B, AP3M1, NRG1, ILK, PFN2, PVALB, SPG7, DAGLA, DNAJC5, CNTNAP1, NAPA, AP3D1 |
| GO:0044463 | Cellular Function | cell projection part | 54, 605 | 0.08617 | AKAP9, RASGRP2, BAIAP2, DCTN2, IGF2BP1, EHD1, CNGA4, PACRG, EPS8, ENKUR, SPATA13, AAK1, TRAK1, CYFIP1, SYT11, KIF13B, TBC1D30, GPR161, AP3M1, NPHP3, BBS9, GRID2, PACSIN1, NRG1, ILK, ITGA5, KCNC3, PDE6B, PFN2, PNOC, SSH1, CFAP46, CCDC40, PRKAR1B, TTYH1, PTH1R, TENM2, PVALB, BBS2, MAP2K4, SLC22A5, SPG7, TIMP2, DAGLA, CA9, JADE1, DNAJC5, MOB2, CNTNAP1, NAPA, BSN, AP3D1, DHRS3, RAB28 |
| GO:0030424 | Cellular<br>Function | Axon | 26, 605 | 0.08617 | IGF2BP1, DAB2IP, EPHB2, AAK1, TRAK1, CYFIP1, SYT11, KIF13B, LDLRAP1, AP3M1, NRG1, ILK, KCNC3, NF1, PFN2, SLC17A7, SEMA6A, PVALB, MAP2K4, SPG7, DAGLA, DNAJC5, CNTNAP1, NAPA, BSN, AP3D1 |
| GO:0098793 | Cellular<br>Function | Presynapse | 24, 605 | 0.08617 | BAIAP2, AAK1, CYFIP1, SYT11, DMXL2, AMPH, GRIK4, GRM8, PACSIN1, HIP1, APBA2, ILK, KCNC3, NF1, SYT17, PFN2, PNOC, PI4K2A, SLC17A7, PVALB, DNAJC5, NAPA, BSN, AP3D1 |

### Relationship Between Early-Onset MD DNAm Markers and PGC MD-Associated Genetic Loci

A total of 6 differentially methylated sites (including DMP/DMRs; *p*=0.002; 95% CI = 2-11) (Supplemental Table 4) and 12 variably methylated sites (including VMPs/VMRs; 95% CI = 5-19, *p*=0.008) (Supplemental Table 5) overlapped PGC GWAS findings. These enrichment results were largely driven by overlap observed with the PGC GWAS locus on chromosome 6 at 27.738-32.848 Mb (Figure 1). At this locus, 5 of 6 differentially and 10 of 12 variably methylated sites overlapped.

**Figure 1.**
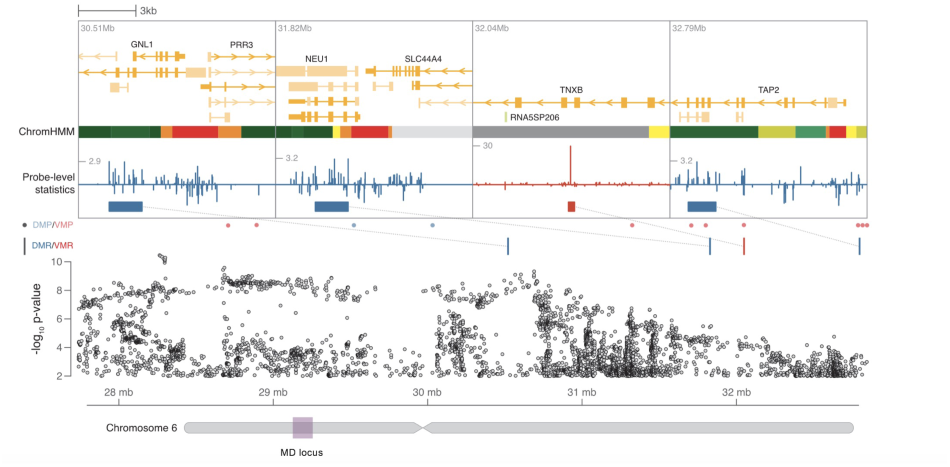
Overlap with PGC GWAS of major depression. The ‘MD locus’ (purple box) represents a region of chromosome 6 ectending from 27.7-32.8 Mb found to be significantly associated by Wray et al. (2018). Summary statistics from this study are plotted for the relevent regional markers in the manhattan plot. Colored ticks represent 3 DMRs (blue) and 1 VMR (red) located in this region. Individual plots above provide a zoomed-in view of the genomic context surrounding each methylation region and probelevel tests statistics. Chromation states within GM12878 lymphoblastoid cells are indicated by color coding the ChromHMM track.

## Discussion

The main objective of the current study was to identify differentially and variably methylated loci and regions that distinguish MZ twins with and without a history of early-onset MD. sis, there was consistency in the functional attributes of the genes related to neural structures and processes as a key differentiating feature between MD affected and unaffected twins. BP gene referenced homophilic cell adhesion and cell-cell adhesion processes, which are involved in neural development and plasticity. A number of cadherins and protocadherins emerged in the cell adhesion gene sets with MD affected twins demonstrating increased variation in two cadherins (*CDH3*, *CDH6*), the clustered protocadherin alpha family (*PCHDA1 - PCDHA13*), the C-type isoforms of PCDHA (*PCDHAC1, PCDHAC2*), and two non-clustered Protocadherins (*PCDH10, PCDH20*). CDHs/PCDHs are a group of calcium dependent cell-cell adhesion molecules that are abundantly expressed in the nervous system and play a major role in multiple steps Across biological process (BP) and cellular function (CF) domains of the gene enrichment analy‐ ontologies essential to neurodevelopment including dendrite arborization, axon outgrowth and targeting, synaptogenesis, and synapse elimination 60–66.

The DNAm profile of the *PCDH*s is known to be responsive to environmental factors^67^, and emerging evidence suggests a role for *PCDH*s in multiple psychiatric phenotypes (e.g., schizophrenia, bipolar disorder, autism) including MD^68,69^. Two small twin studies observed that several cadherin/protocadherin genes demonstrated differences in DNAm between twin pairs repeatedly ^70^ as well as a history of MD or an anxiety disorder^71^. A related study observed increased DNAm in *PCDH* gene families with the highest enrichment of hypermethylated sites in the *PCDHA* genes located in the hippocampus of suicide completers with a history of severe childhood abuse^19^. At the genetic variant level, a recent meta-analysis with over 89,000 MD cases detected a SNP (rs9540720) in the non-clustered *PCDH9* gene to be significantly associated with MD case status at the genome-wide level^72^, and a related gene that encodes a protein of the same family (*PCDH17)* was found to confer risk for mood disorders^73^. Moreover, expression patterns of *Pcdh* genes have been examined in rodent brain regions in volved in the neural circuity of MD^74^, with results indicating high expression levels in subregions of the hippocampus and basolateral amygdaloid complex^74,75^. Moreover, *Pcdh* gene expression was reduced in both structures following electroconvulsive shock (ECS), which is interesting given that ECS is an effective treatment for chronic, intractable MD symptomatology^76^ and supports involvement of *PCDHs* in neural plasticity. The clustered *PCDHA* family also is strongly expressed in serotonergic neurons^77–79^ and *PcdhαC2* is necessary for axonal tiling and assembly of serotonergic circuitries^86^. Thus, a deeper understanding of *how* the serotonergic neural network is formed in the young developing brain may be critical to identifying etiological determinants of MD and other psychiatric conditions.

The Histone Deacetylase 4 (*HDAC4*) gene was altered in MD affected twins, exhibiting increased variation and emerging as part of a genomic region displaying higher mean levels com‐ discordant for elevated depression symptoms pared to MD unaffected twins^80^. *HDAC4* is highly expressed in the human brain,^81^ and the sub‐ cellular localization of the *HDAC4* protein is purportedly modulated by *N*-methyl-D-aspartate (NMDA) receptors, a specific type of ionotropic glutamate receptor ^82^. In humans, increased *HDAC4* expression is associated with memory deficits^80,82^, which is consistent with the cognitive processing difficulties observed with MD^83,84^. Moreover, drugs that inhibit *HDAC4* have antidepressant-like effects on behavior^85–88^. Similarly, Neuroregulin1 (*NRG1*) exhibited increased variation and emerged in the gene enrichment analysis as a gene involved in multiple cellular functions. *NRG1* promotes myelination in the central nervous system (CNS) and evidence suggests an association between *NRG1* and cognitive deficits linked to psychiatric disorders^89,89–95^.

The overlap between our DNAm results and variants identified in the latest PGC GWAS of depression were examined to better understand the possible role of early-onset MD DNAm events among these genetic associations. A total of 6 differentially methylated and 12 variably methylated sites overlapped PGC findings. These enrichment results were primarily driven by overlap observed with the PGC GWAS locus on chromosome 6, which falls in the extended MHC region. This region of the genome also was the most significant association in the PGC’s GWAS of schizophrenia^96^ and a GWAS that included a combined sample of schizophrenia and bipolar disorder cases^97^. The MHC is densely populated with genes related to neuronal signaling and plays a role in immune functioning. MHC findings are consistent with evidence of an immune component involved in the pathophysiology of psychiatric conditions.

A number of additional genes/gene regions previously linked to MD (e.g., *CX3CR1,CACNA1A*, *CDC42BPB*)^99,102-104^. For example, serotonin related genes, *HTR5A* and *HTR5A-AS1*, exhibited greater variance in MD affected twins. This gene is a member of the 5HT (serotonin) receptor family, which has been implicated in MD and other psychiatric conditions^105–107^. Related, corticotropin releasing hormone receptor 2 (*CRHR2*) and cholinergic receptor nicotinic beta 2 subunit (*CHRNB*2) demonstrated increased variance in MD affected twins. Thus, complex genetic important for controlling syn‐ aptic plasticity and memory function disorders such as MD reflect a large number of independent genetic factors that each contribute a small amount of variance to disease susceptibility and multiple psychiatric diseases likely share genetic risk factors.

Limitations associated with the current study design include its reliance on an all Caucasian sample, which diminishes generalizability of findings to other races. The current study also relied on peripheral blood tissue for DNAm. Although brain tissue is preferred to study the pathophysiology of MD, evidence suggests moderate consistency in DNAm between blood and brain tissues^15,71^. Consistent with epidemiological findings^108^, females represented the majority of the sample. Thus, increased numbers of males are needed to determine potential sex-related differences. The cross-sectional design of the current study also does not allow for determination of epigenetic differences as cause or consequence of MD onset. Strengths of the current study de sign include its use of the MZ co-twin control method in conjunction with a sensitive developmental window to reveal genome-wide DNAm biomarkers associated with early-onset MD.

## Acknowledgements

The research outcomes presented in this manuscript were supported by a NARSAD Independent Investigator Award from the *Brain and Behavior Research Foundation* to the first author (RRN). The development of analytic methods were supported by a NARSAD Independent Investigator Award from the *Brain and Behavior Research Foundation* to the last author (TPY).

## Supplemental Methods

The primary definition of MD affected status required presence of at least one DSM-5 MDE lasting at least two weeks^109^. MD was considered present if criteria were met at either a definite or a probable level. A definite diagnosis was made when a twin endorsed at least five DSM-5 Criterion A MDE symptoms (e.g., low energy, recurrent thoughts of death, inability to concentrate) of which at least one symptom had to include the presence of low mood and/or anhedonia most of the day, nearly every day for at least two weeks during their worst episode. Probable diagnoses were made when a twin endorsed at least 4 symptoms, one of which had to include the presence of low mood and/or anhedonia most of the day, nearly every day. The use of definite and probable levels allows for the inclusion of twins who may not be currently symptomatic and, therefore, have to rely on retrospective memory to determine their symptom presentation. The use of probable levels of diagnosis also is justified given that twins expressing nearly all symptoms of a MDE are more similar to a case than a control symptomatically and for depressive risk factors^110^. This approach to diagnosis was introduced as part of the Research Diagnostic Criteria and has been applied in many studies^111–114^. To qualify for MD at the definite or probable level, participants also had to report that their depression symptoms caused clinically significant distress or functional impairment (i.e., Criterion C). MD unaffected status was defined as no lifetime diagnosis of MD at the probable or definite threshold. The expanded MD section of the CIDI-SF followed by the lifetime MD algorithm is presented below.

## Lifetime Major Depression Diagnostic Questions

### Criterion A (MDE Symptoms)

1. Have you ever had a time in your life when you felt sad, blue, or depressed or two weeks or more in a row? (yes/no)
2. Have you ever had a time in your life lasting two weeks or more when you lost interest in most things like hobbies, work, or activities that usually give you pleasure? (yes/no)
3. During those worst two weeks, did the feelings of sadness or loss of interest usually last all day long, most of the day, about half of the day, or less than half of the day? (All day long, Most of the day, About half of the day, Less than half of the day)
4. Did you feel this way every day, almost every day, or less often during the two weeks? (Every day, Almost every day, Less often)

If a participant responded “yes” to questions 1 OR 2 AND endorsed “all day long” or “most of the day” to question 3 AND responded “every day” or “almost every day” to question 4, they entered the MD section and were queried regarding other DSM-5 Criterion A symptoms of MD (see below).

5. Thinking about those same two weeks, did you feel more tired out or low on energy than is usual for you? (yes/no)
6. Did you gain or lose weight without trying or did you stay about the same? (Gained weight, Lost weight, Gained and lost weight, stayed about the same, I was on a diet)
7. About how much weight did you gain/you lose/your weight change? (numeric response)
8. Did you have more trouble falling asleep or staying asleep than you usually do during those two weeks? (yes/no)
  a. Did that happen every night, nearly every night, or less often during those two weeks? (Every night, Nearly every night, Less often)
9. During those two weeks, did you have a lot more trouble concentrating or making decisions than usual? (yes/no)
10. People sometimes feel down on themselves, no good, or worthless, or have excessive guilt and blame themselves for things. During that two-week period, did you feel this way? (yes/no)
11. Did you think a lot about death – either your own, someone else’s, or death in general during those two weeks? (yes/no)

### Criterion C (Functional Interference)

12. Did you ever tell a professional about these problems (such as a medical doctor, psychologist, social worker, counselor, nurse, clergy, or other helping professional)? (yes/no)
13. Did you ever take medication for these problems? (yes/no)
14. How much did these problems interfere with your life or activities – a lot, some, a little, or not at all? (A lot, Some, A little, Not at all) ^*^additional questions regarding age of onset, timing of last episode, etc. were queried.

## MD Algorithm

Questions 6 and 7 were recoded so that a response of “Gained weight” or “Gained and lost weight,” along with at least a response of 5 pounds in question 7, was coded as endorsement of weight gain/loss. Participants who endorsed question 8 also were presented with question 8a.Sleep symptoms were considered present if a participant answered “Every night” or “Nearly every night” on question 8a. Responses to queries 1, 2, 5, weight gain, sleep symptoms, 9, 10, and 11 were summed as the total number of depression symptoms endorsed during the twin’s worst MDE. Participants were deemed to have experienced significant distress if they endorsed question 14 as “a lot” or “some” and/or if they endorsed question 12 or 13 as “yes”. MD at the full threshold level was coded positive if the participant endorsed question 1 or 2 along with question 3 as “All day long” or “Most of the day,” and question 4 as “Every day” or “Almost every day,” and at least 4 other depression symptoms for a total of at least 5 symptoms. MD at the full threshold level also required significant distress. MD was coded positive at the probable level if the participant endorsed question 1 or 2 as well as question 3 (“All day long” / “Most of the day”) and question 4 (“Every day” / “Almost every day”) and at least 3 other depression symptoms for a total of 4 symptoms. The probable level also required endorsement of significant distress (i.e., endorsed question 12, 13, or 14 as “a lot” or “some”).

## Exclusionary Criteria

Participants were not eligible for the current study if they met any of the following criteria: 1) current use of psychotropic medications (e.g., antianxiety/antidepressants) or medications with psychotropic effects (e.g., beta-adrenergic blockers), b) diagnosis of an autism spectrum disorder, c) diagnosis of an intellectual disability, d) diagnosis of a spatial learning disorder, or prior testing indicating an IQ below 70, e) seizure without a clear and resolved etiology, f) current or past episodes of psychosis, g) serious, not stabilized illness (e.g., liver, kidney, gastrointestinal, respiratory, cardiovascular, endocrinologic, neurologic, immunologic, or blood disease), h) inadequate production of human growth hormone, i) sensory integration disorder, j) congenital adrenal hyperplasia, k) adrenal inefficiency, l) deaf with bicochlear implants, m) cancer (current or past diagnosis), and n) pregnancy (current or lifetime). If only one twin from the twin pair met any of the exclusionary criteria, the whole pair was excluded.

## Zygosity

Zygosity status (monozygotic [MZ] versus dizygotic [DZ]) for adolescent twins (age ≤ 17) was determined based on parent-report about physical similarities between twins. Adult twins (age ≥ 18) not accompanied by a parent/legal guardian completed the zygosity questionnaire about themselves. Prior research has demonstrated high validity for this zygosity assessment as compared to blood^115^ and DNA evaluations of zygosity^116^. MZ zygosity based on questionnaire assessment was confirmed using 65 SNP control probes included on the Infinium HumanMethylation450 (450K) array to verify sample identity. The control probes target polymorphic sequences, and values for each probe cluster into three groups corresponding to genotype. Together, the control probes provide strong support for validating zygosity. Only data from twins confirmed to be MZ were included in analyses.

## Genome-wide DNAm Measurement and Processing

Genomic DNA was isolated from whole blood according to standard methods using the Puregene DNA Isolation Kit (Qiagen). An aliquot of 1 microgram DNA per subject was processed by HudsonAlpha Institute for Biotechnology for bisulfite conversion (Zymo Research EZ Methylation Kit) and genome-wide methylation assayed on the Infinium Human Methylation 450K Bead-Chip microarray, which interrogates 485,764 features. Twin pairs were localized to the same slide to minimize any potential artifactual differences in DNAm patterns due to batch effects.

Details of the 450K microarray have been previously described^117^, and raw data processing was performed according to best practices reported in recent publications^50,118^. Intensity values from the scanned arrays were processed using the *minfi* Bioconductor package^119^ in the R programming environment (R Development Core Team 2015). Confirmation of self-reported race was made by sample clustering derived from principal components estimated from ancestry informative probes^120^.

Quality was assessed both quantitatively and visually to identify deviant samples^119^. Beta values were derived as the ratio of the methylated probe intensity to the sum of the methylated and unmethylated probe intensities^121^. Beta value density plots from each array were inspected to tag poor performing arrays based on a large deviation from the rest of the samples. Probes were filtered if they had a detection P-value of greater than 0.01 in at least 10% of samples or if they have been previously identified as cross-hybridizing^122^ leaving a total of 455,828 probes to analyze. Quantile normalization adapted to DNAm arrays^123^ was applied to adjust the distribution of type I and II probes to the final set of screened sample arrays and probes.

For all statistical tests, *beta* values were transformed using the *M*-value procedure to promote normality and calculated as a logit transformation of the methylated and unmethylated intensity ratio along with an added constant to offset potentially small values^121^. Correlations between major experimental factors and the top 10 principal components of *M*-values across all arrays were inspected to identify extraneous structure that may account for any batch effects^124^. ComBat was used to remove average differences across arrays due to slide groupings^125^. Blood cell proportions were inferred for each sample to account for cellular heterogeneity^126^.

---

**Supplemental Table 1.** Statistical contrasts fit to estimate the degrees of association between MD affected/unaffected status and genome-wide DNA methylation markers.

|  | Concord-<br>ant MD T1 | Concordant MD<br>T2 | Discordant<br>T1 | Discord-<br>ant T2 | Concord-<br>ant Healthy<br>T1 | Concordant<br>Healthy T2 |
| --- | --- | --- | --- | --- | --- | --- |
| **contrast 1** | -1 | -1 | -1 | 1 | 1 | 1 |
| **contrast 2** | -1 | -1 | 2 | 0 | 0 | 0 |
| **contrast 3** | 0 | 0 | 0 | 2 | -1 | -1 |
| **contrast 4** | -1 | 1 | 0 | 0 | 0 | 0 |
| **contrast 5** | 0 | 0 | 0 | 0 | -1 | 1 |

**Note: T1 = Twin 1 and T2 = Twin 2.**

**Supplemental Figure 1.**
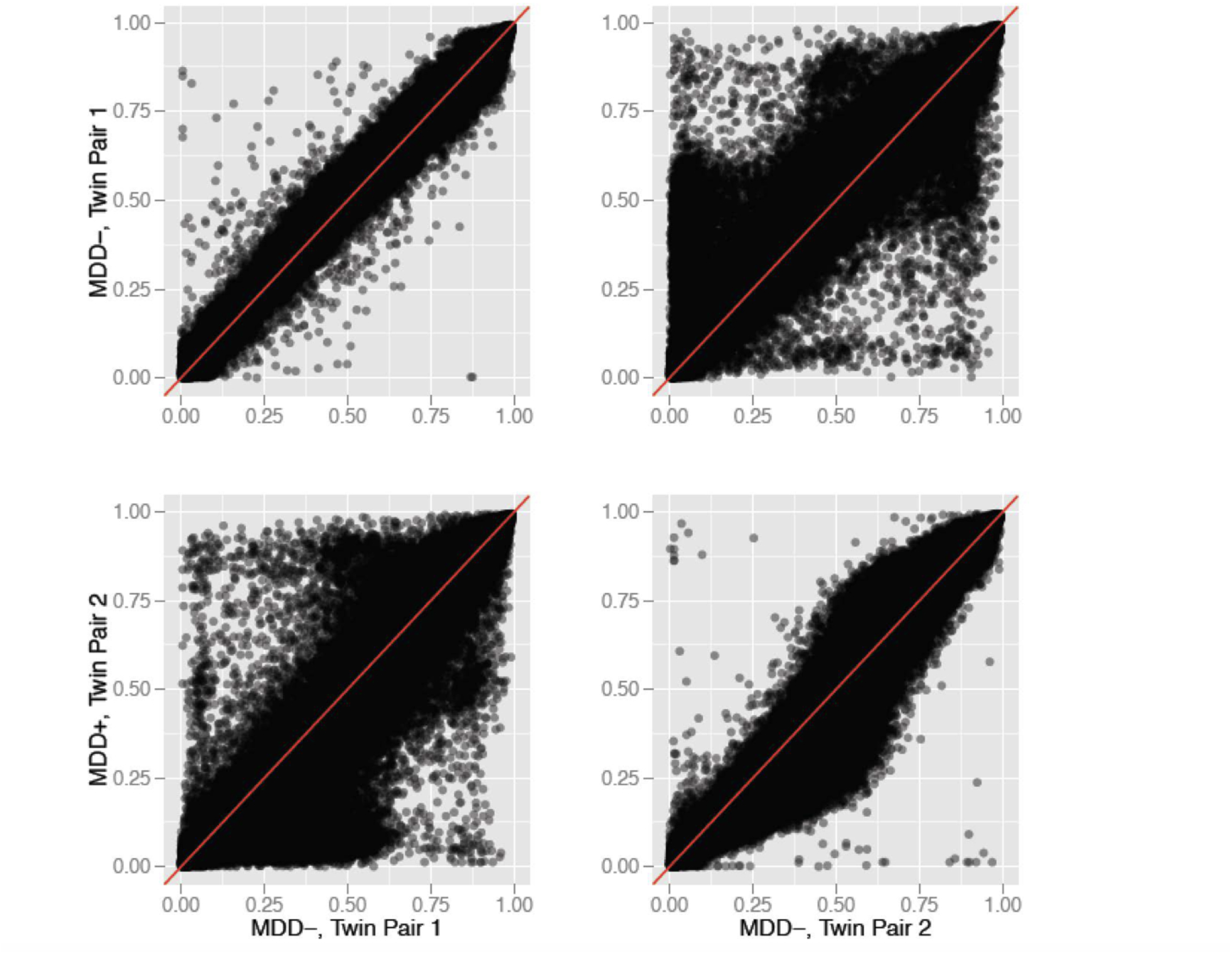
Bivariate distribution of DNAm Probes, where one twin pair is discordant for a history of MD and the other is concordant healthy. A twin member from the discordant pair and the concordant healthy pair is then crossed in the upper right and lower left panels, creating “non-MZ” pairs. This figure shows that the bivariate distribution of DNAm marks differ between the concordant healthy and discordant twins, with the discordant pair showing greater variation between twins. The lower left panel, which represents an unrelated case-control, illustrates the substancial increase in variance, underscoring the utility of the co-twin control design to study environmental risk.

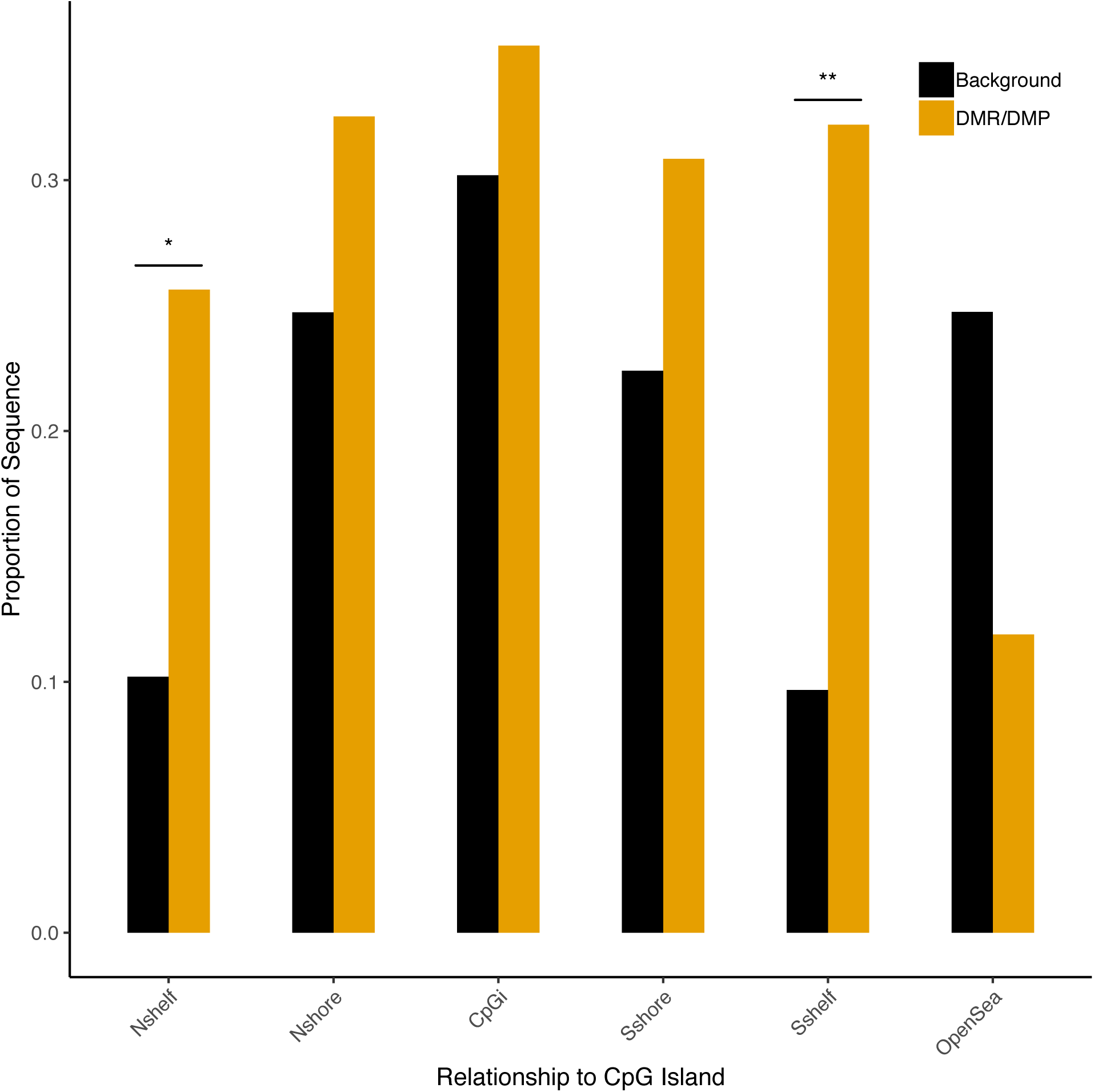

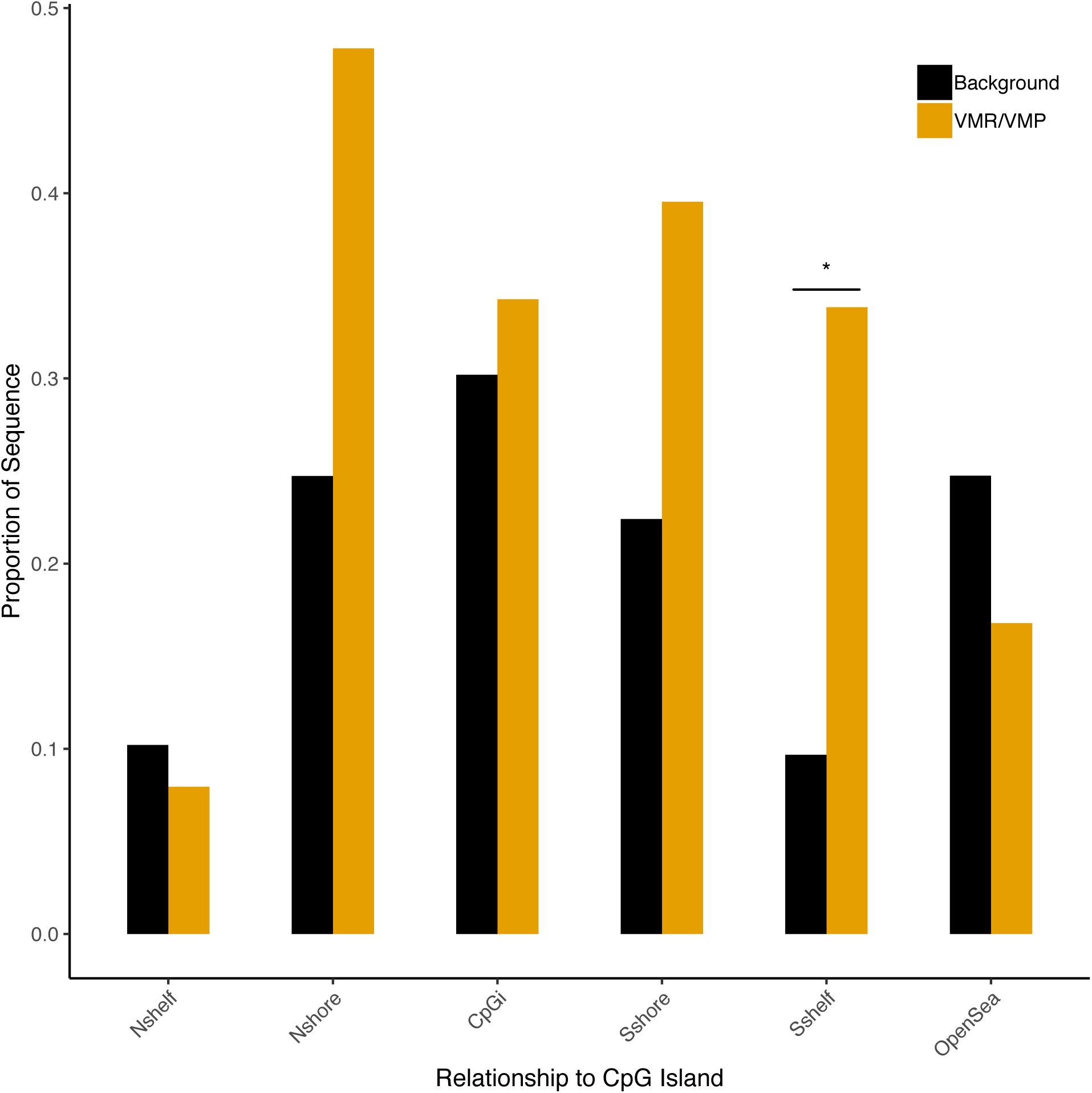

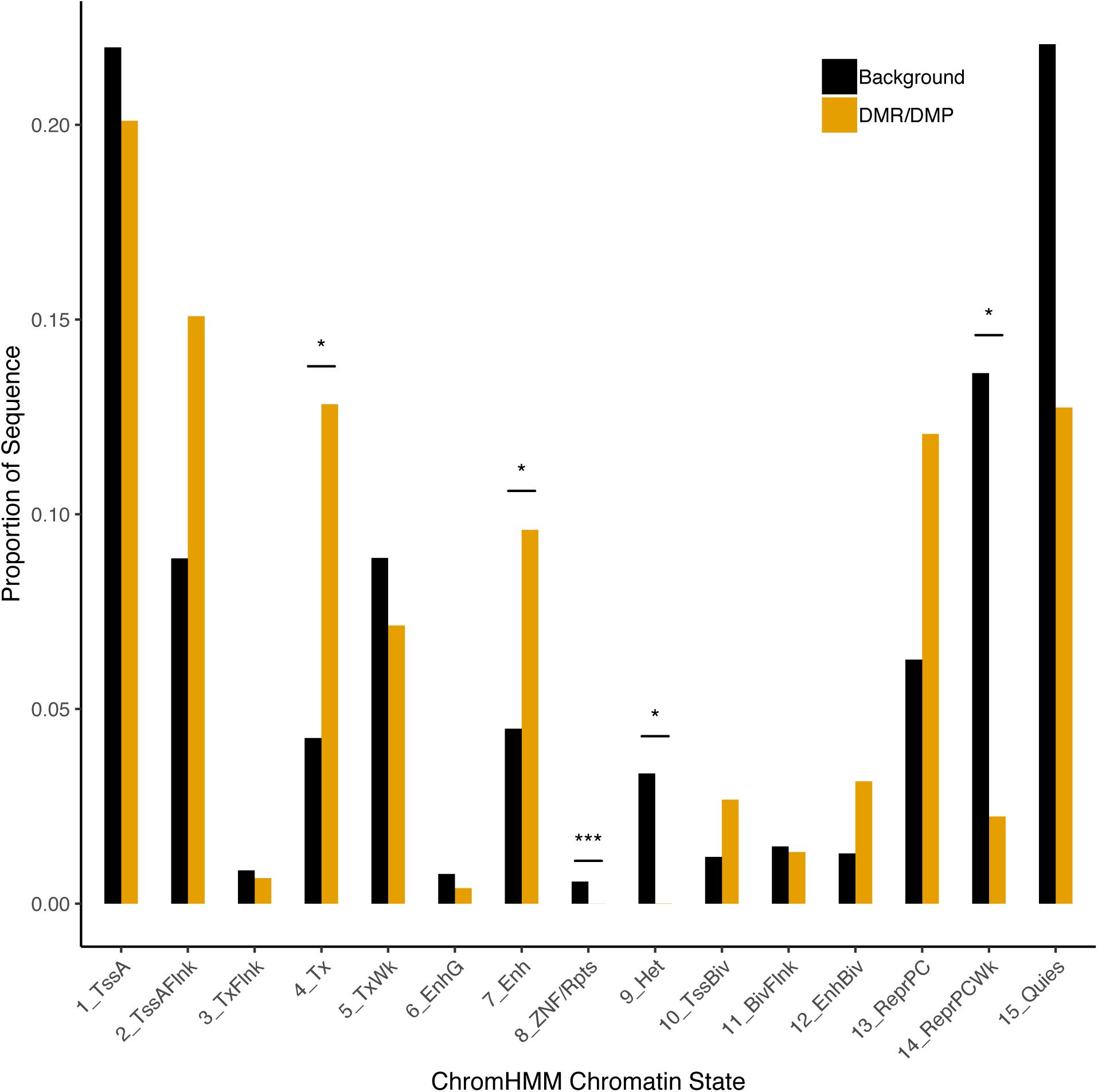

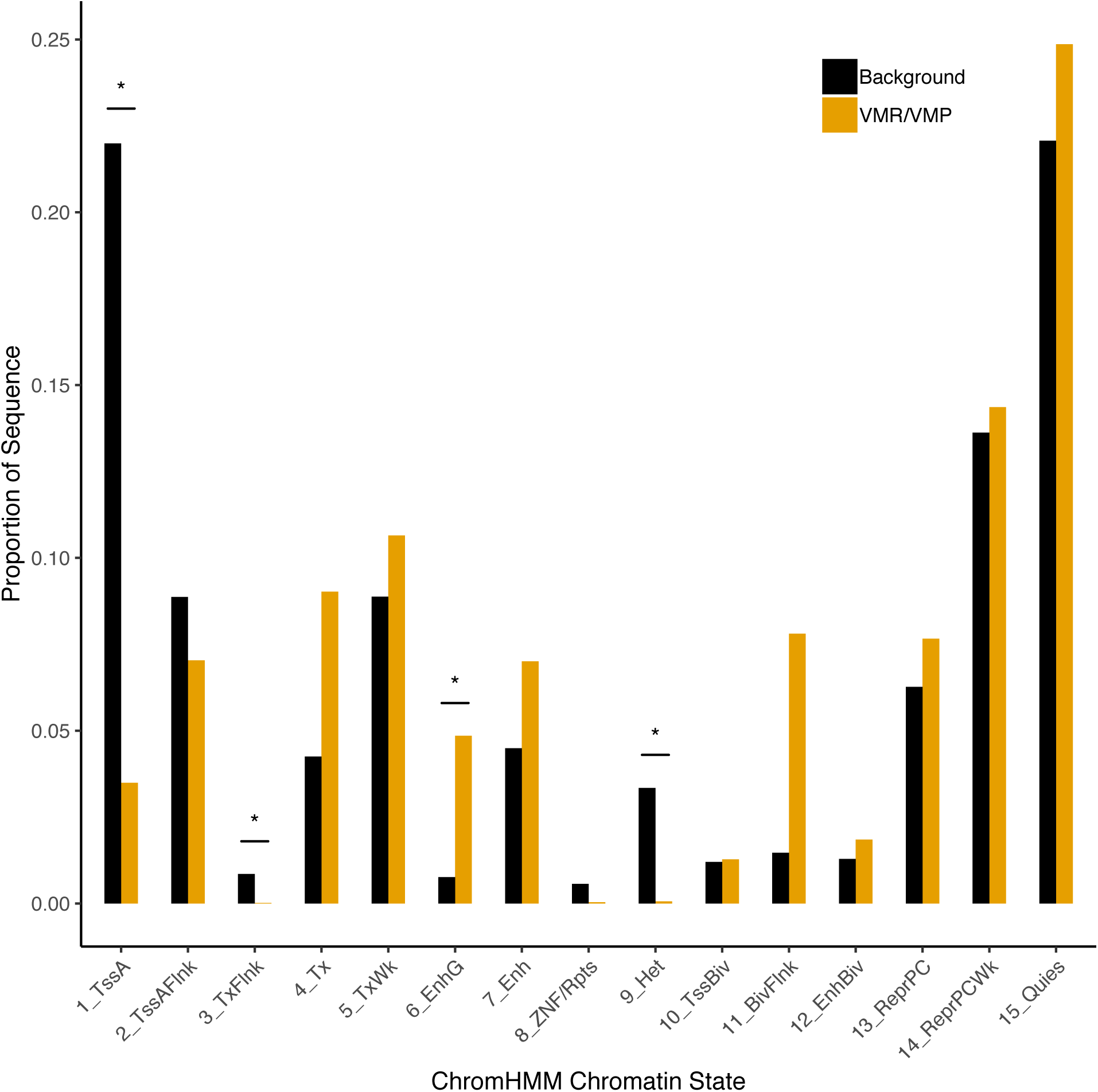

**Supplemental Table 2.**
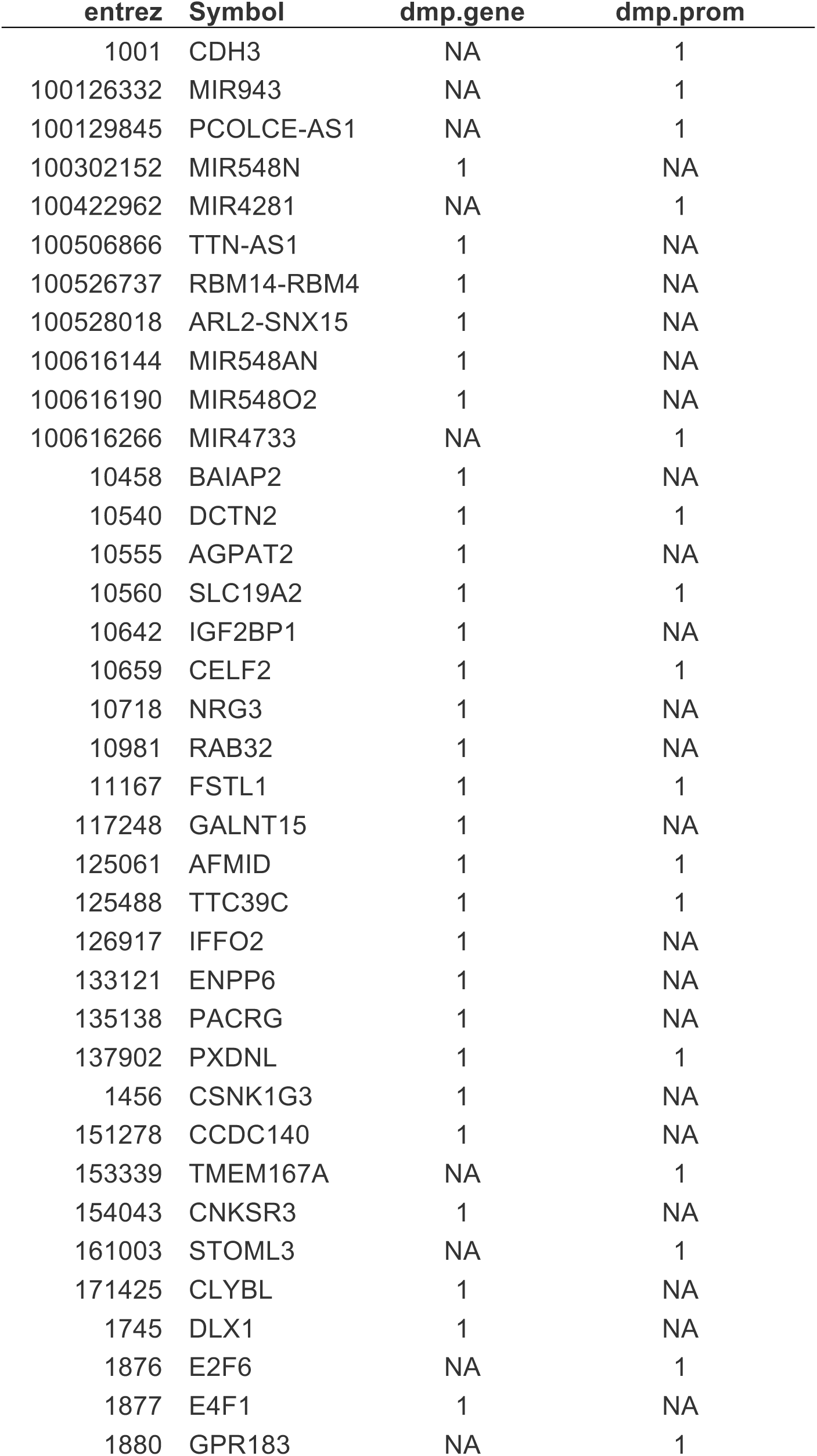
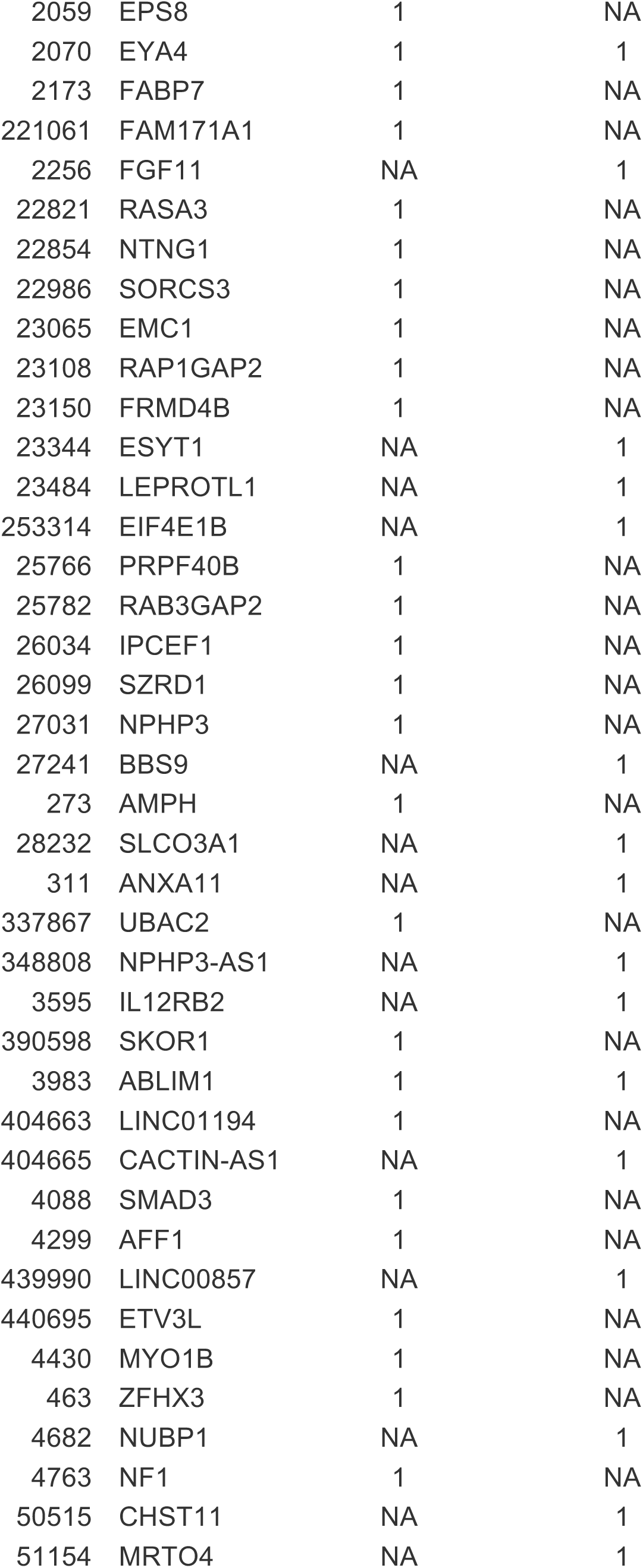
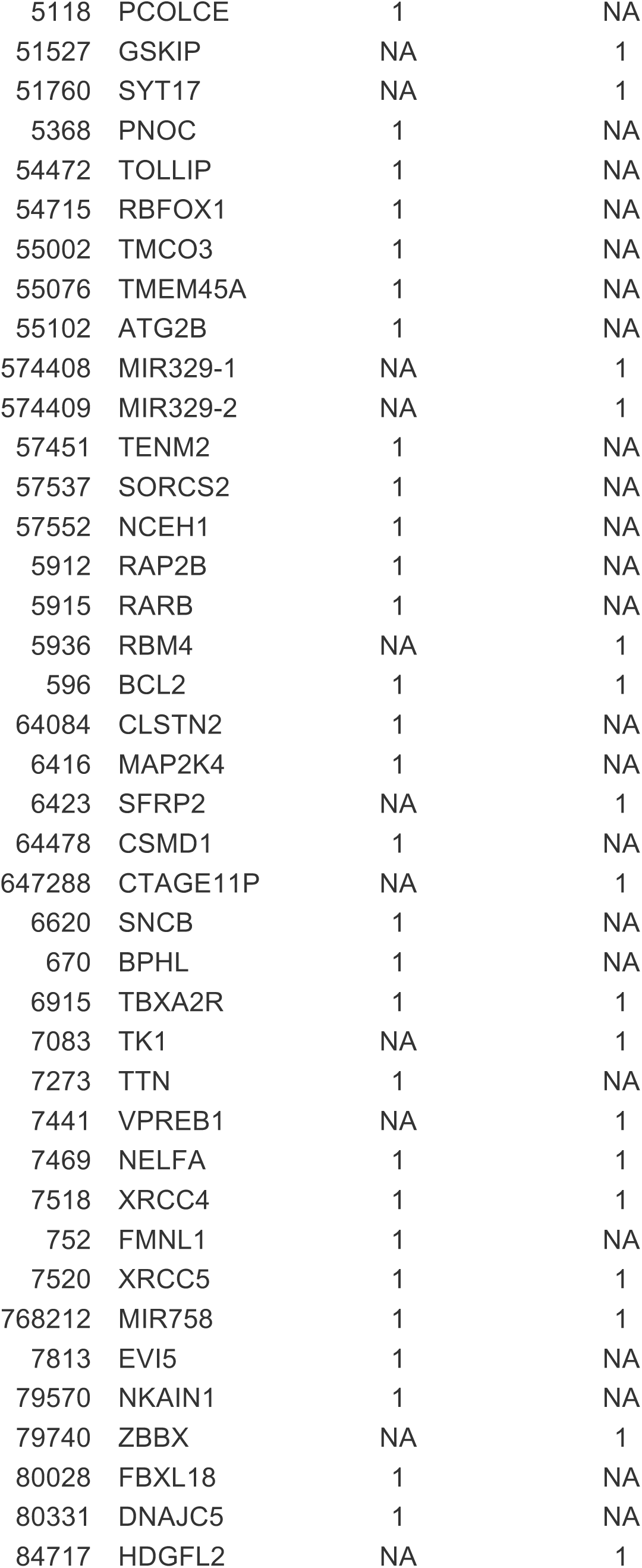
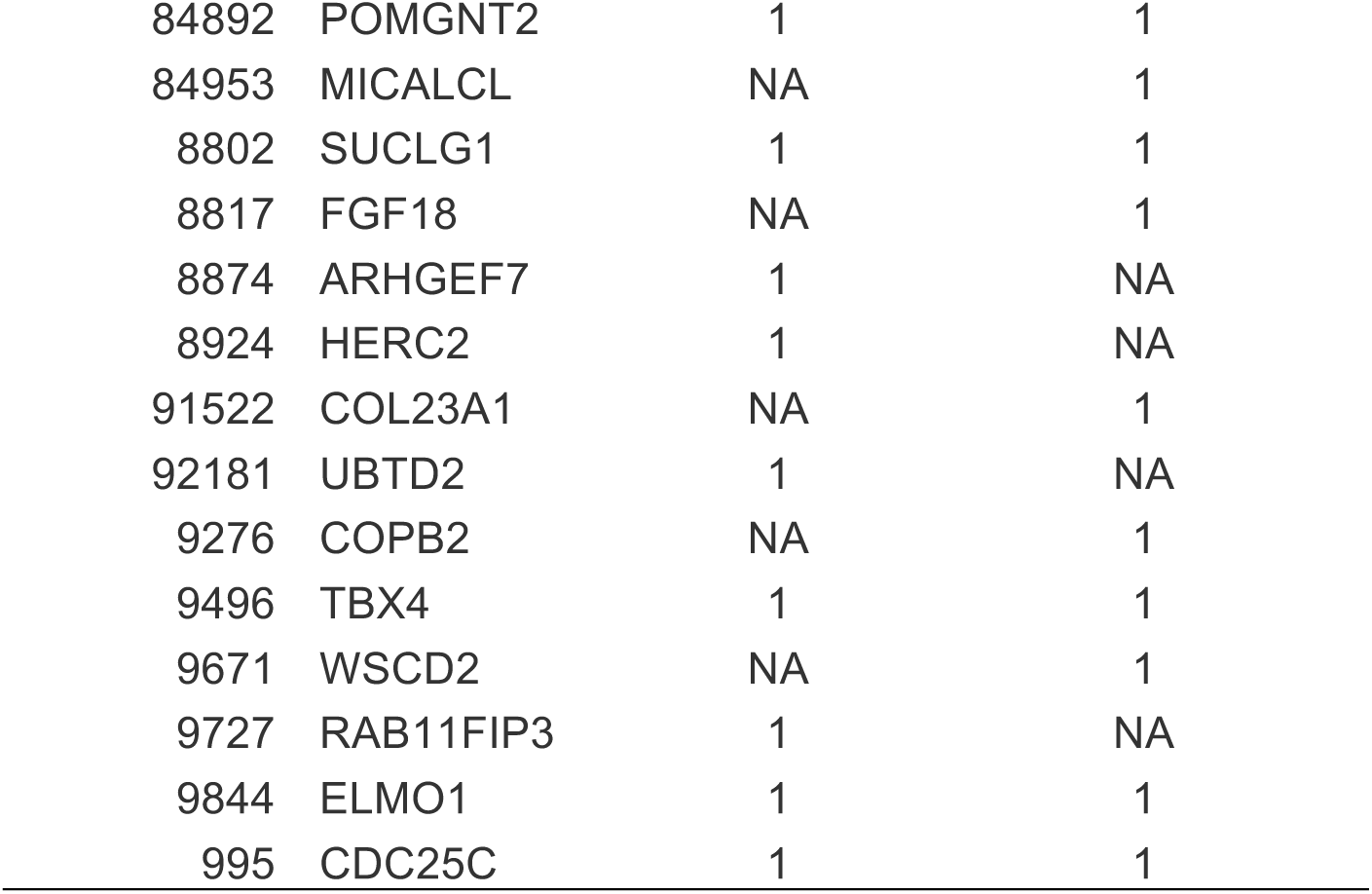
DMP significant gene/promoter hits identified in the MD affected versus MD unaffected contrast.

**Supplemental Table 3.**
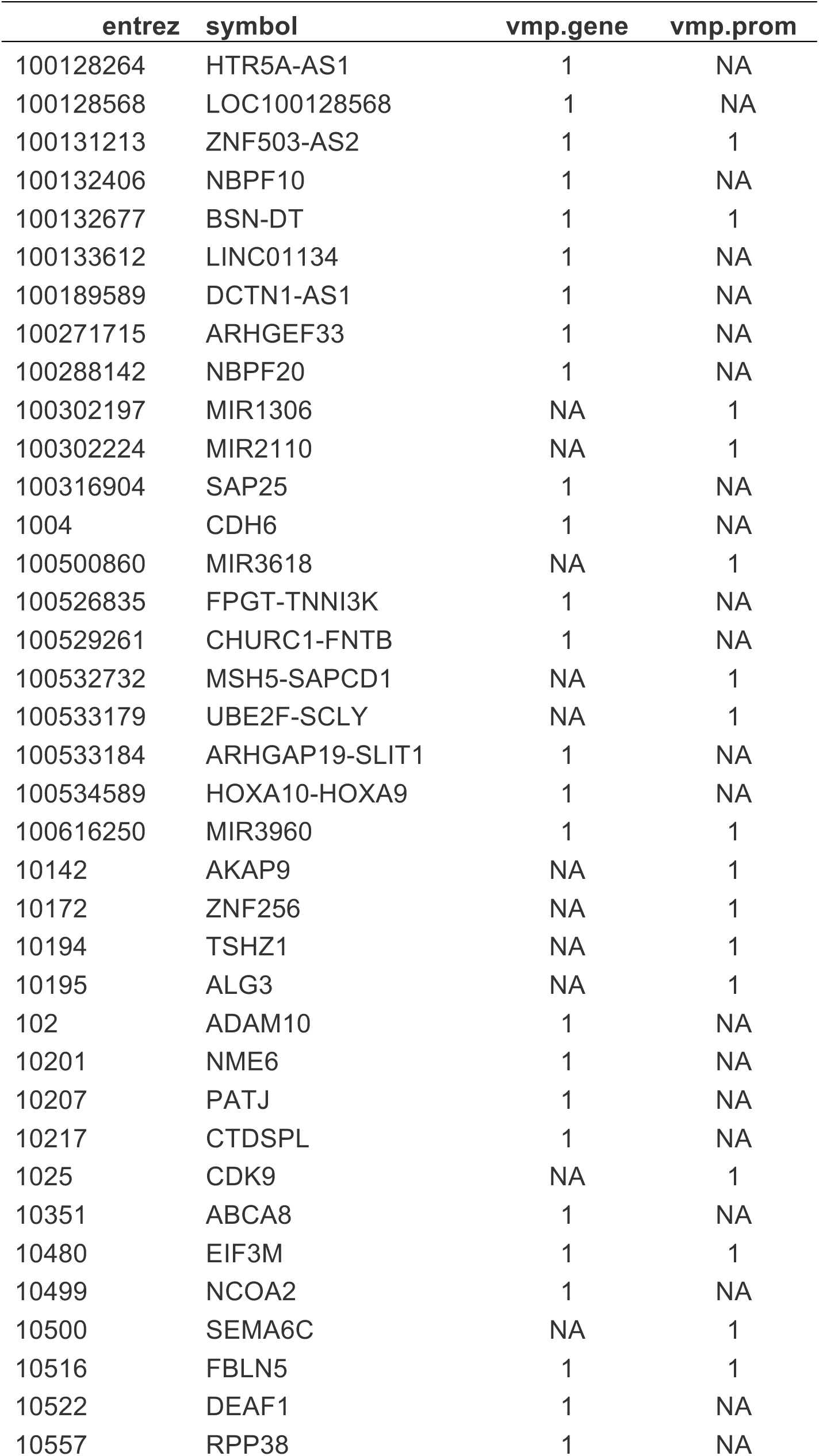
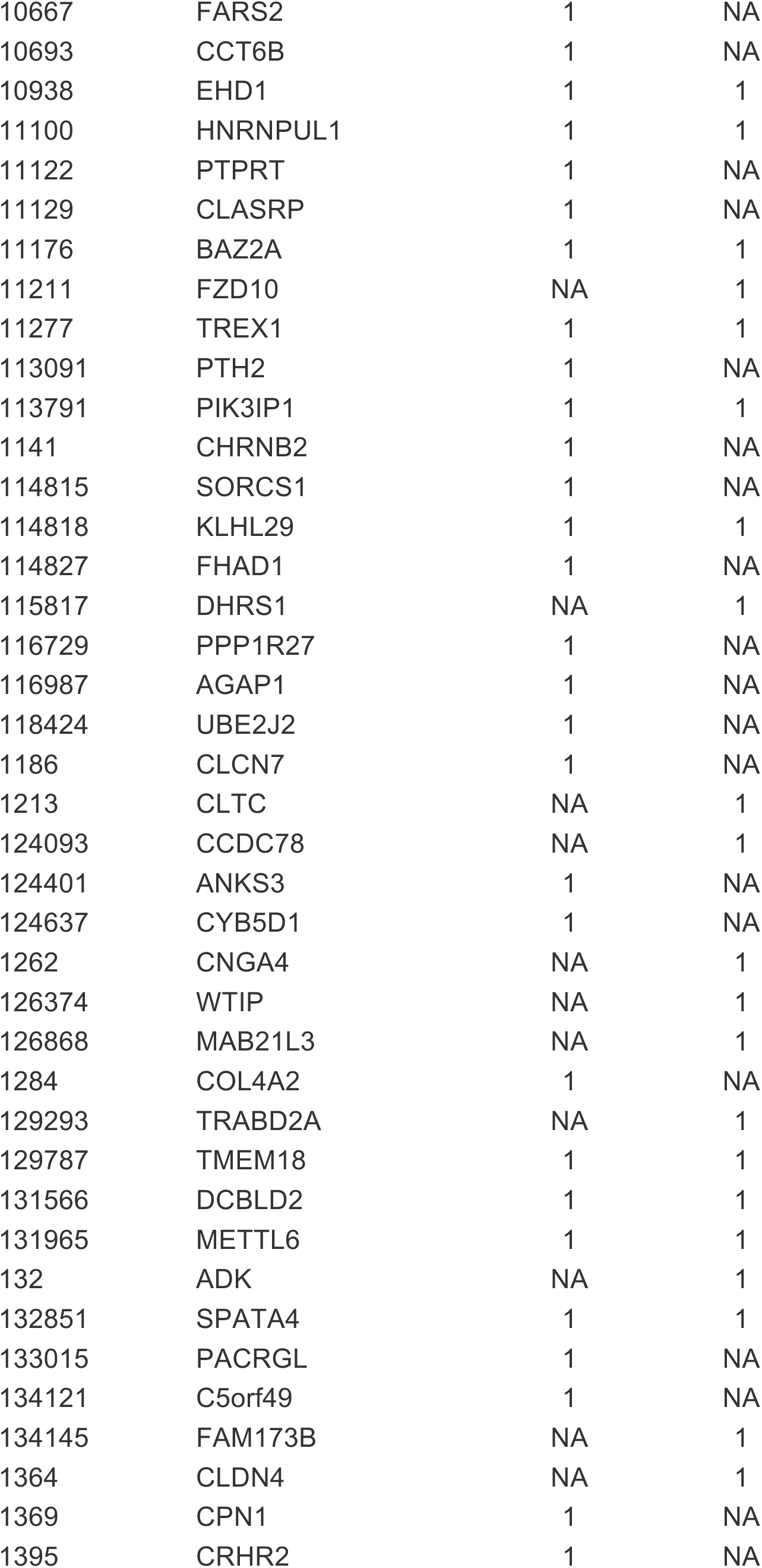
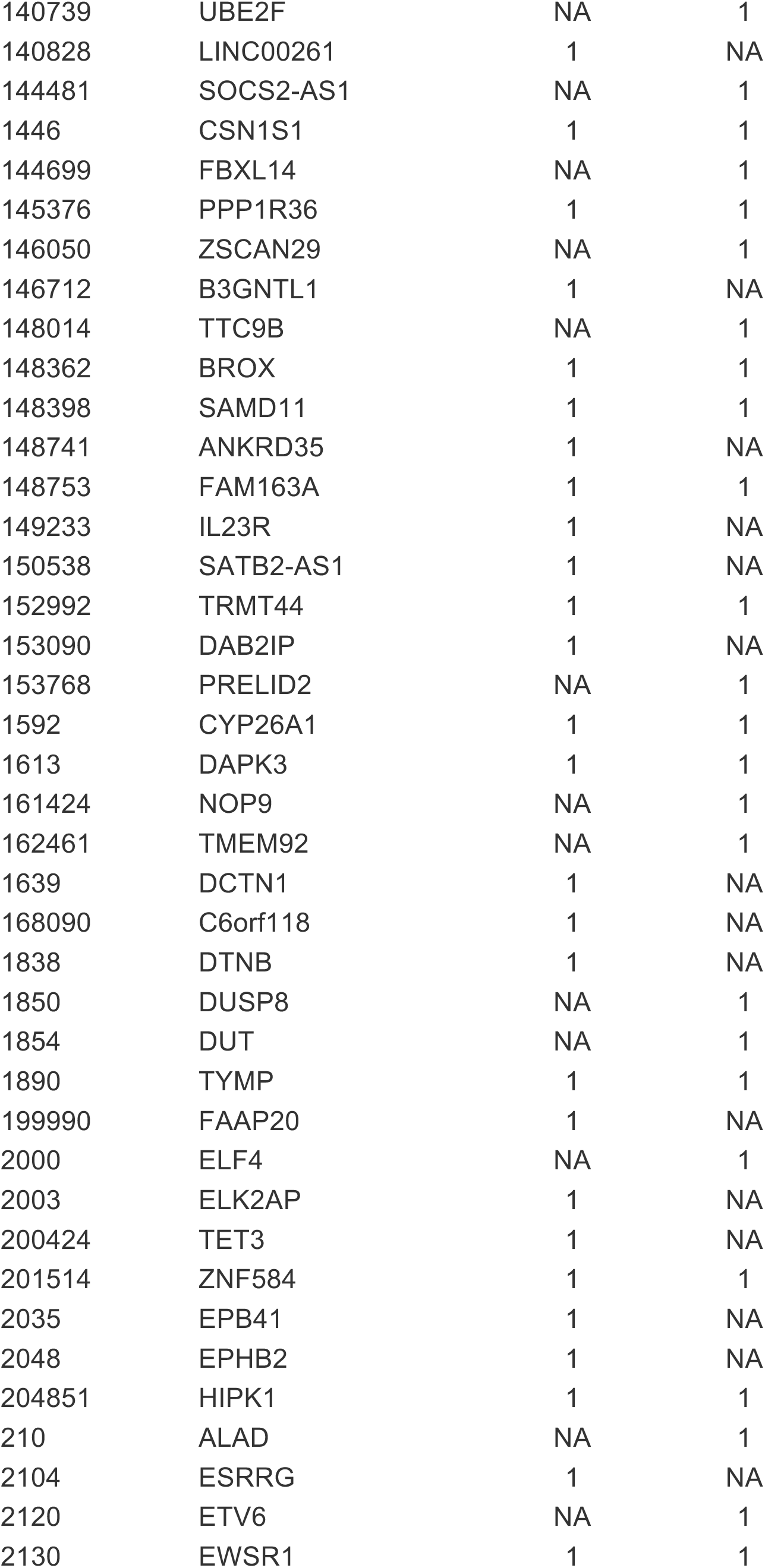
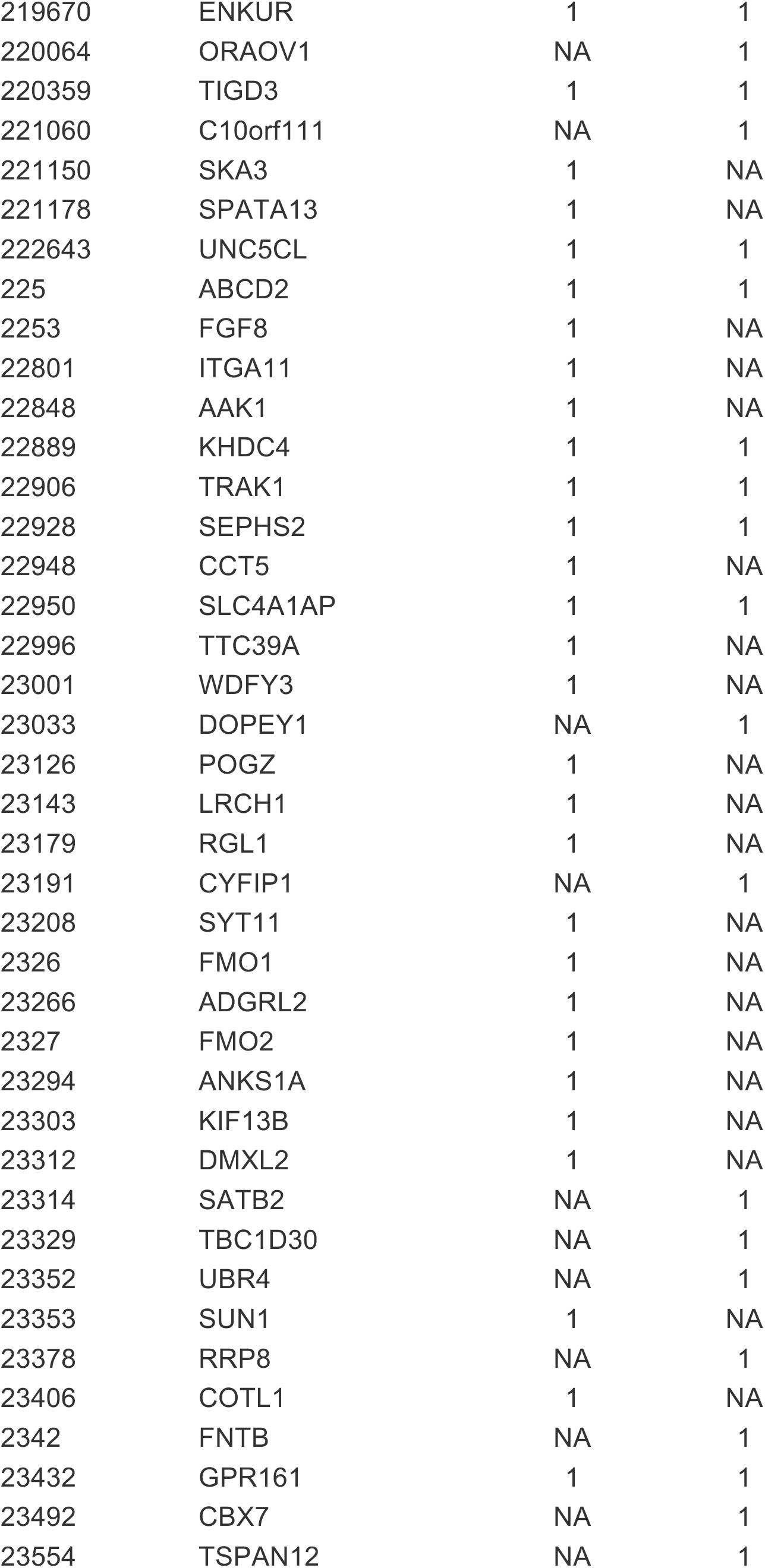
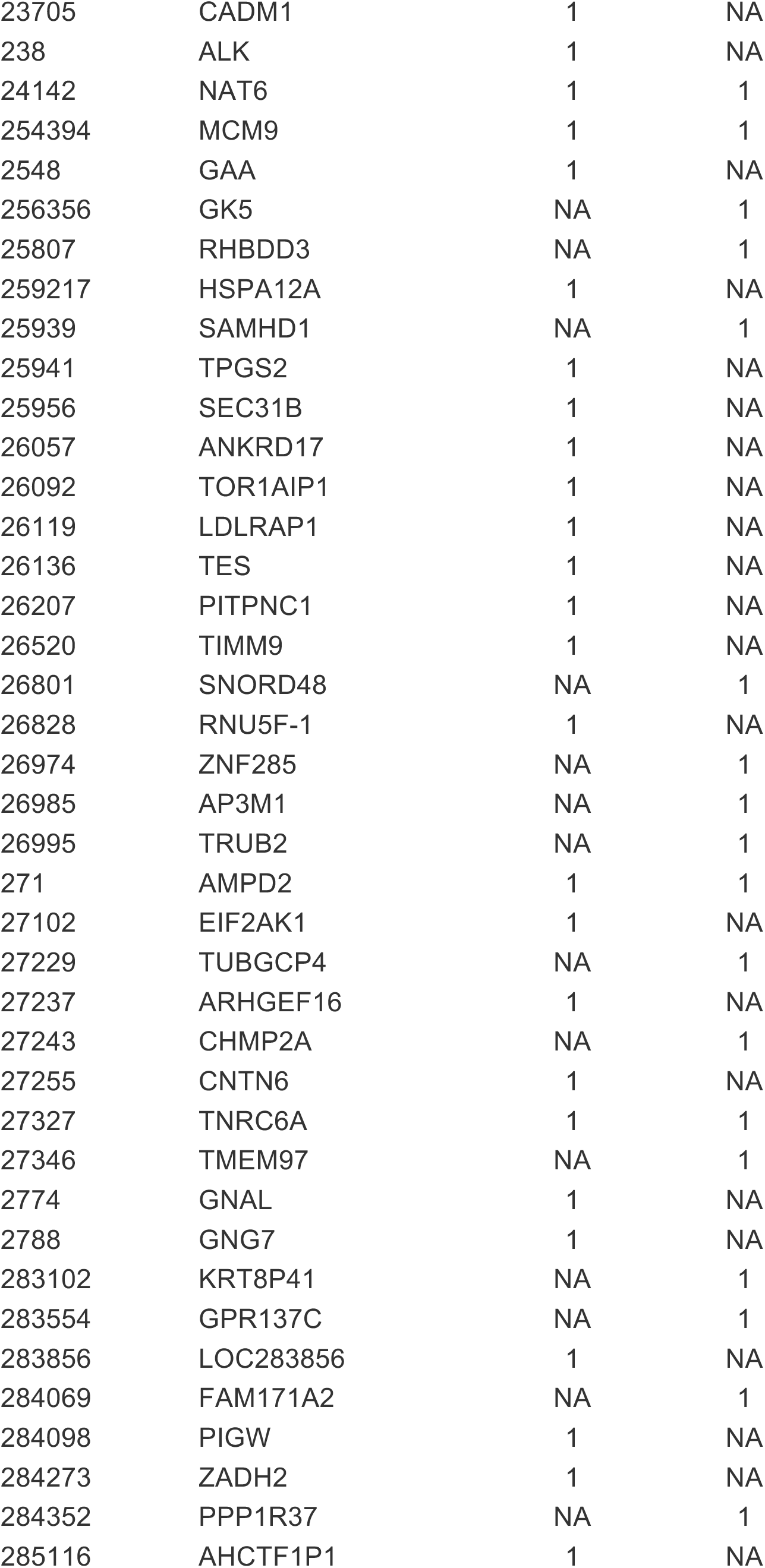
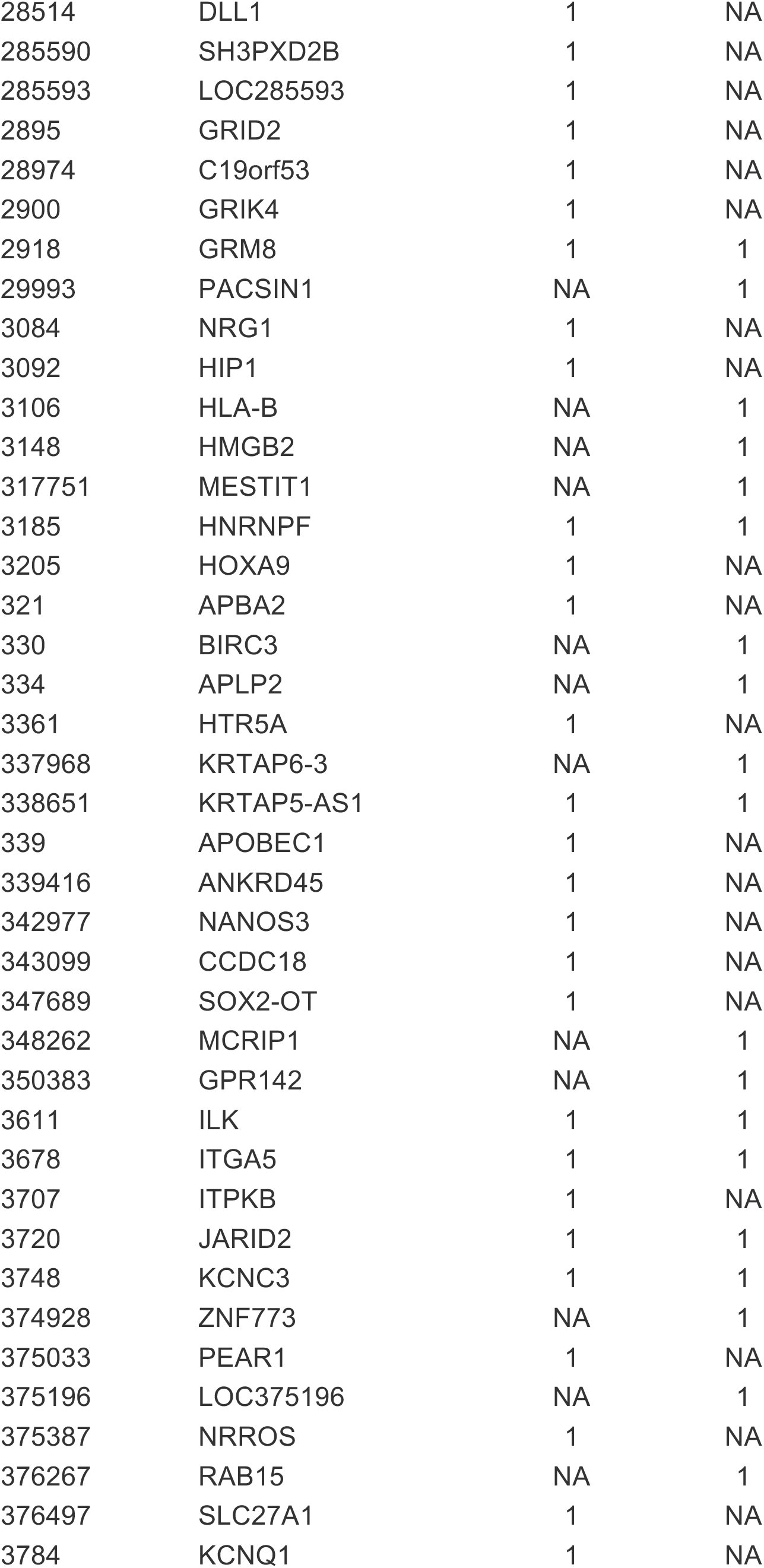
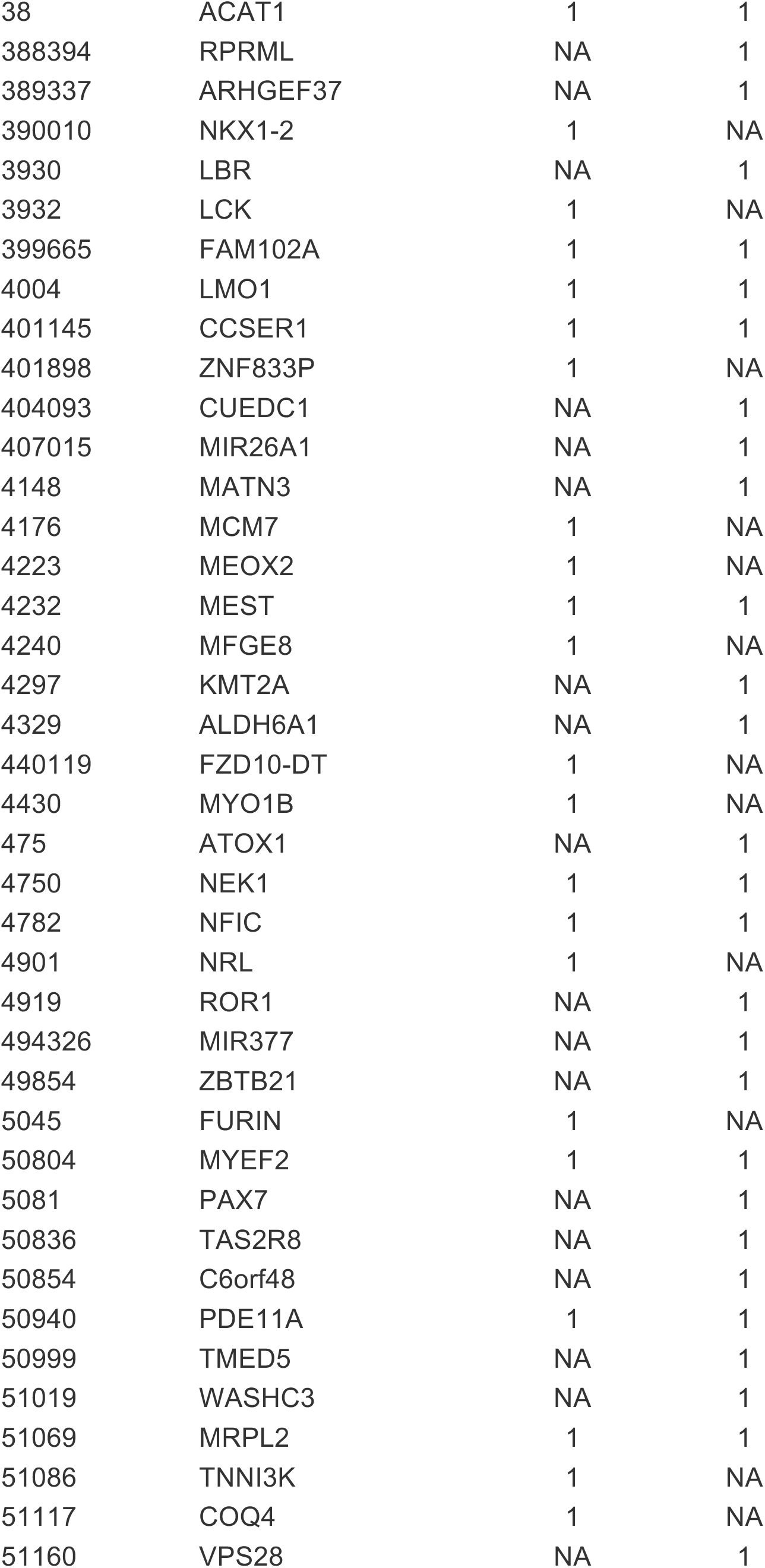
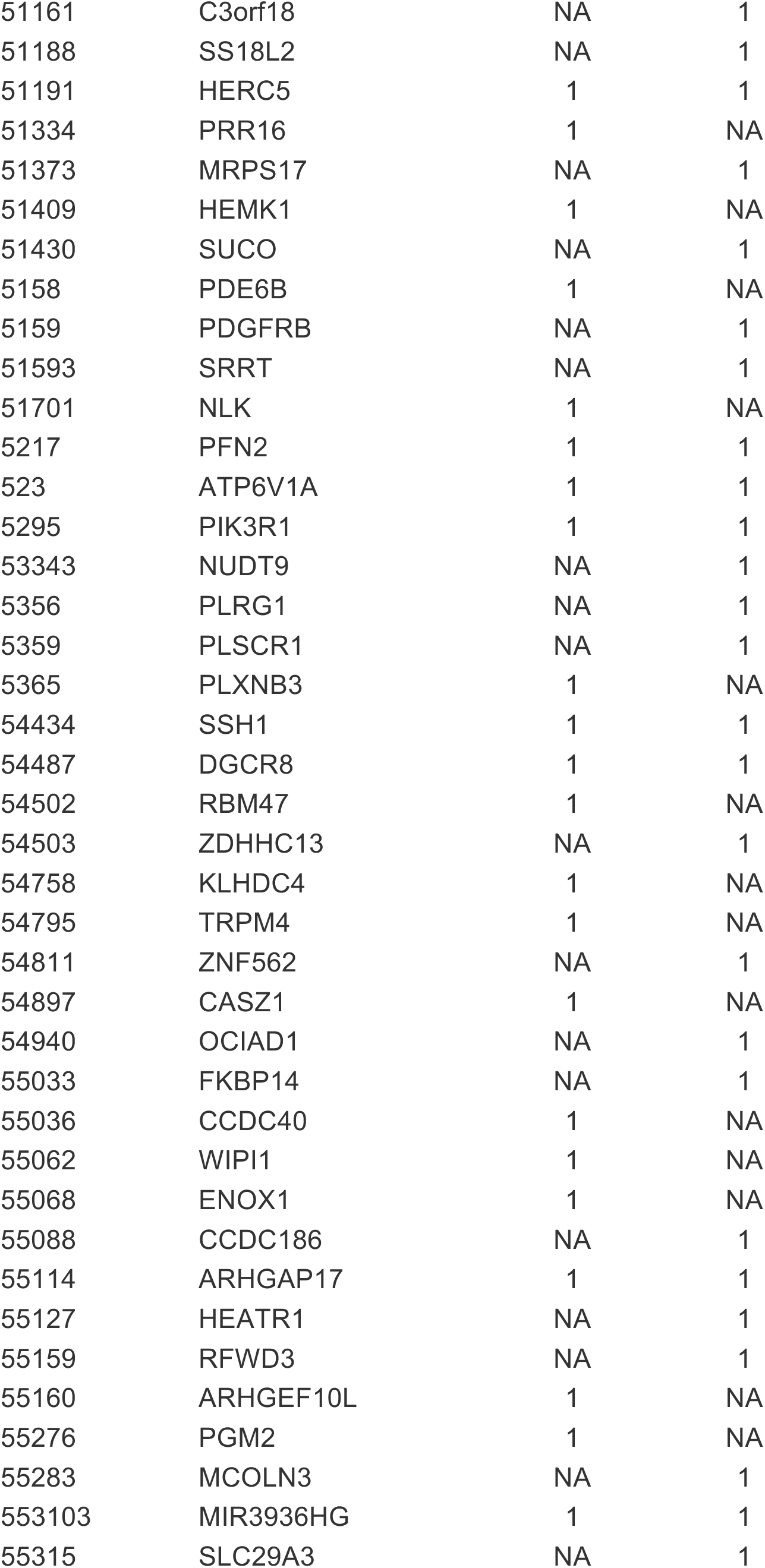
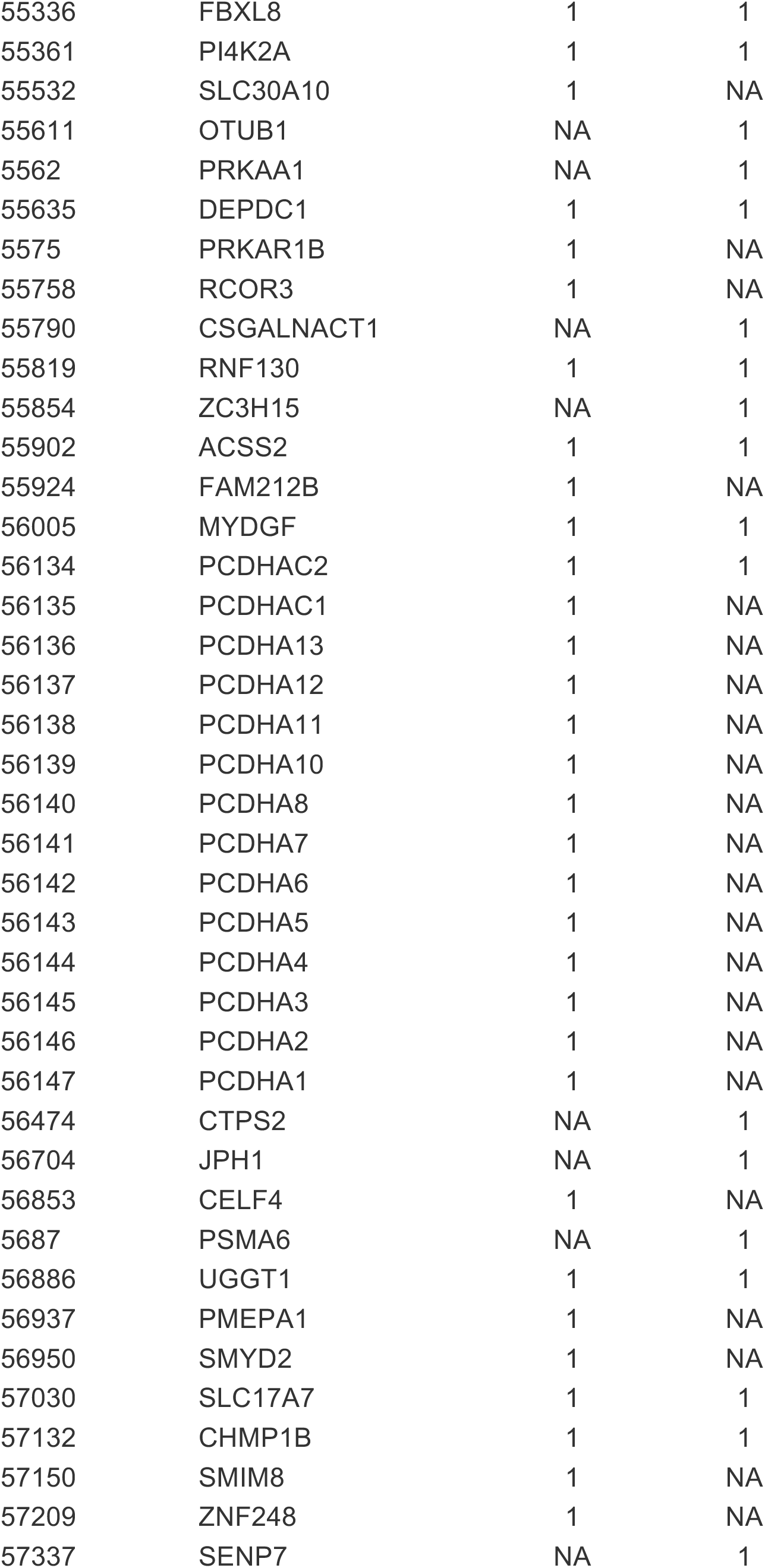
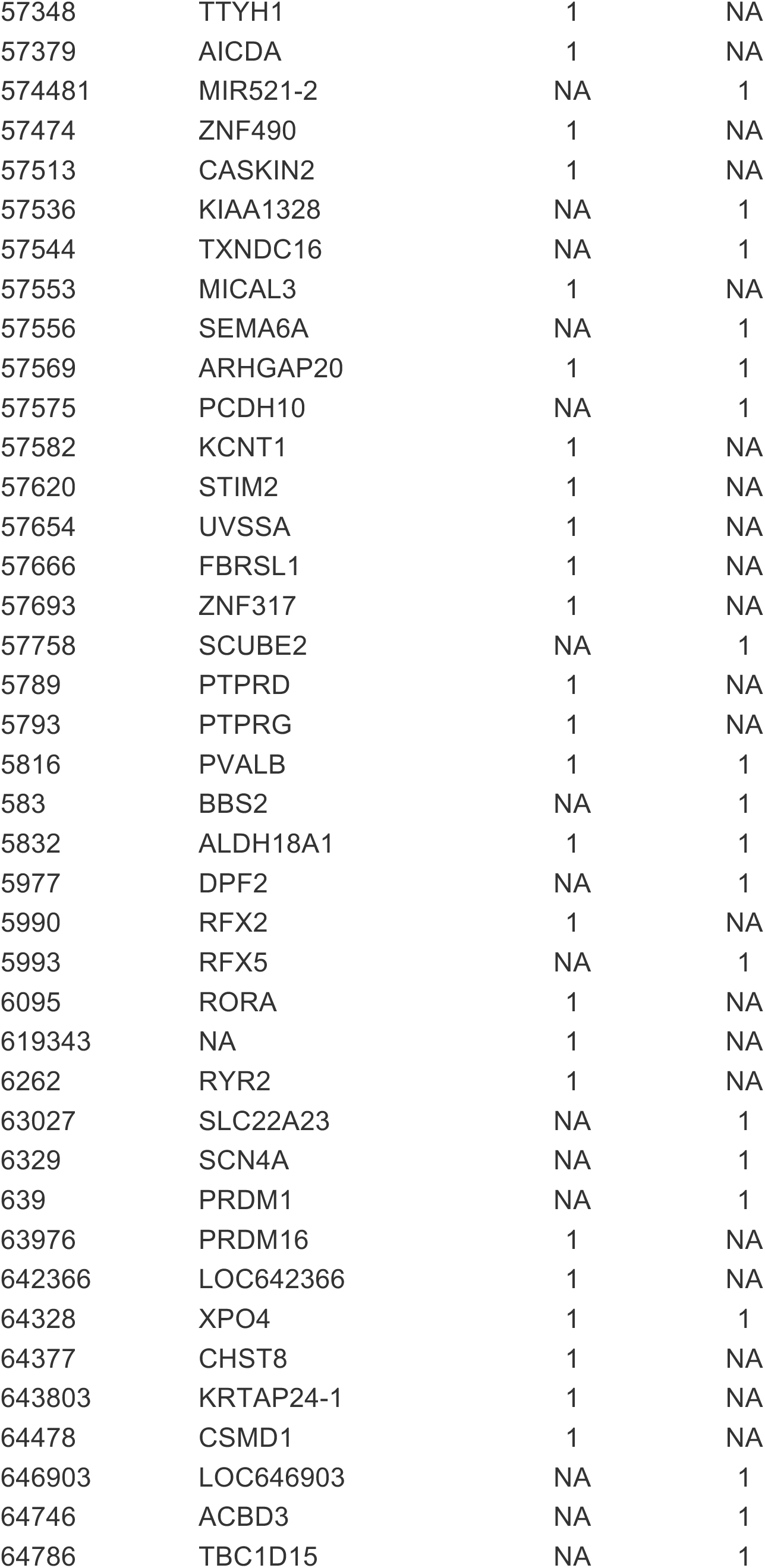
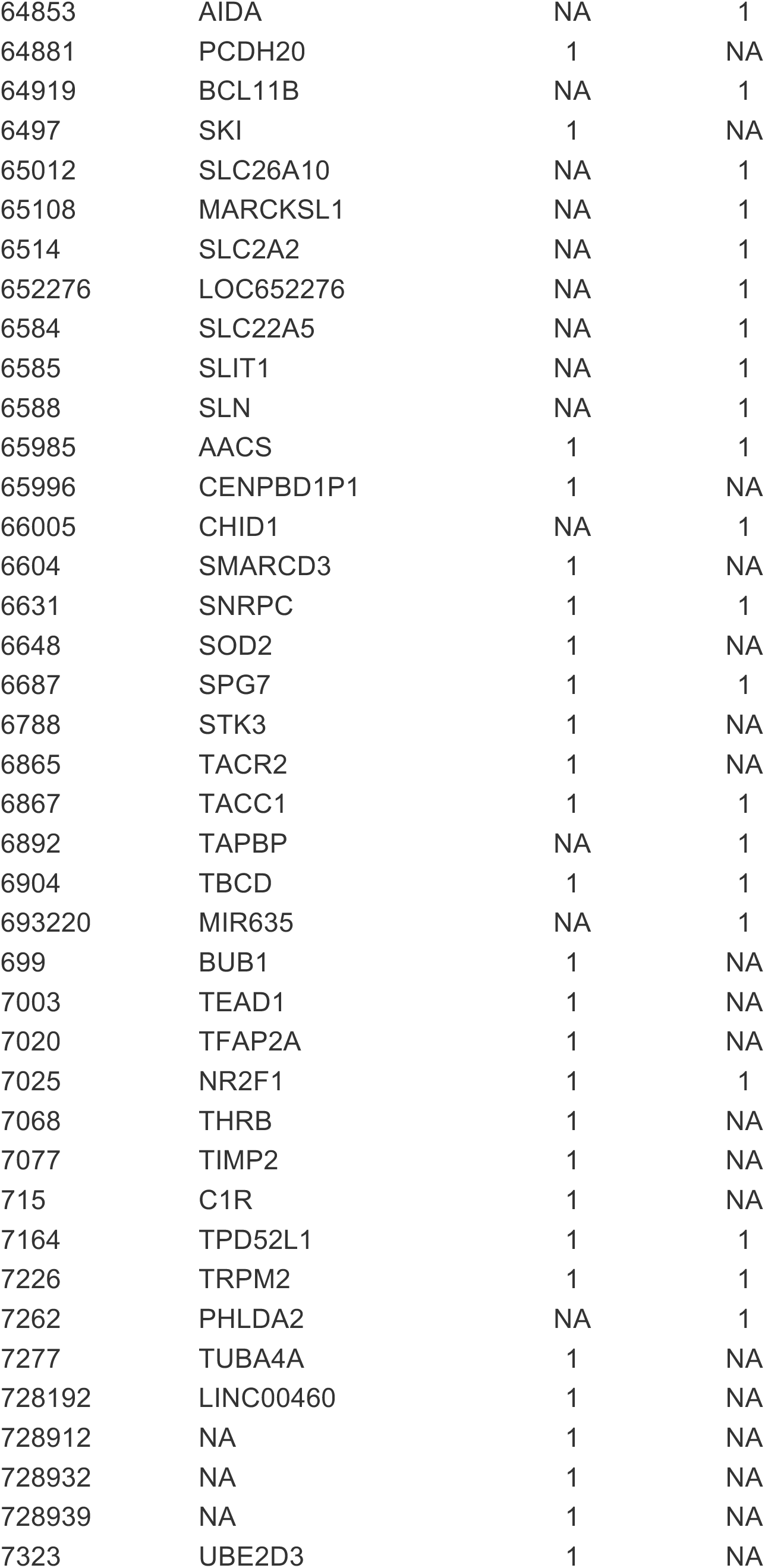
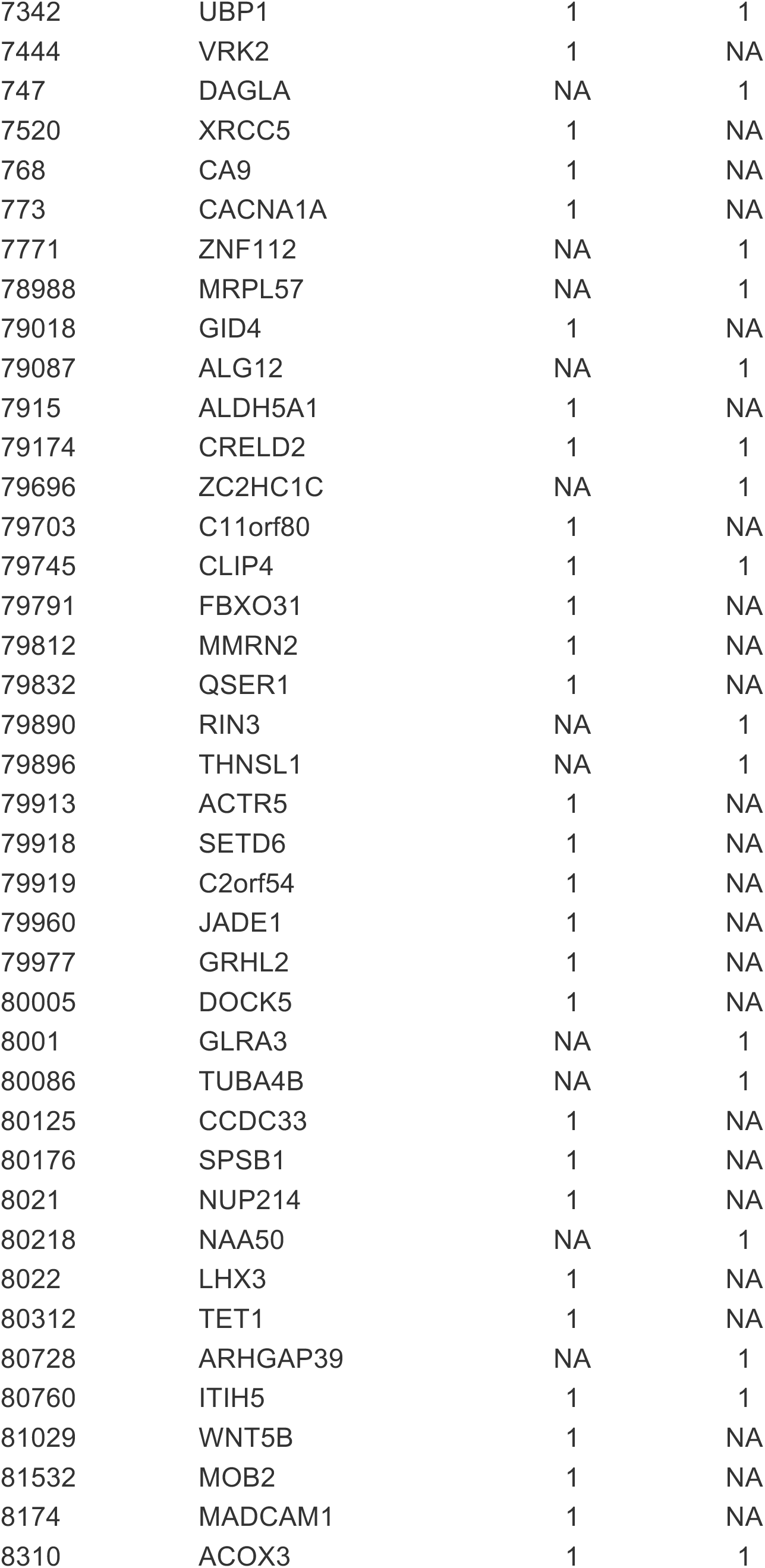
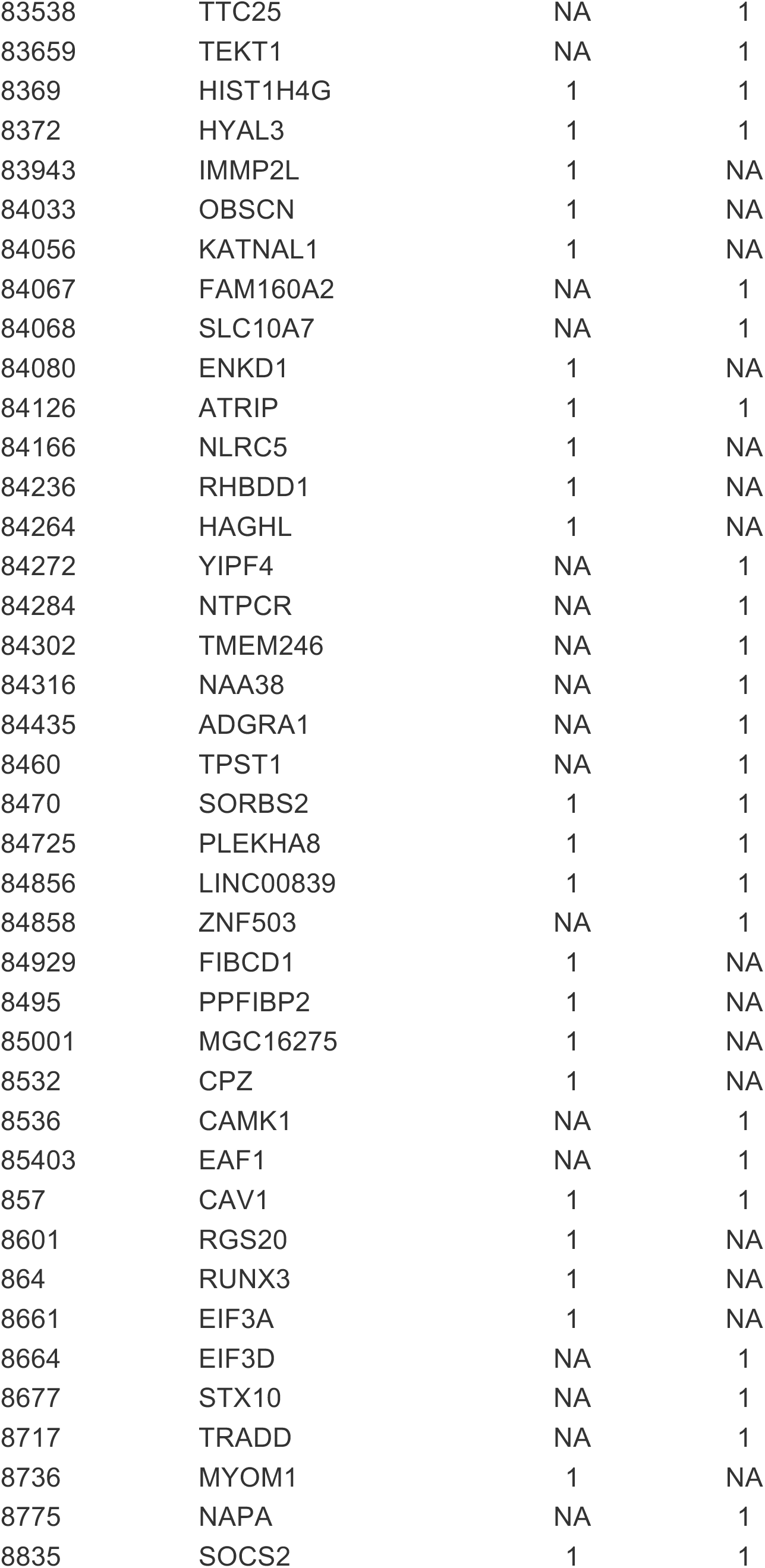
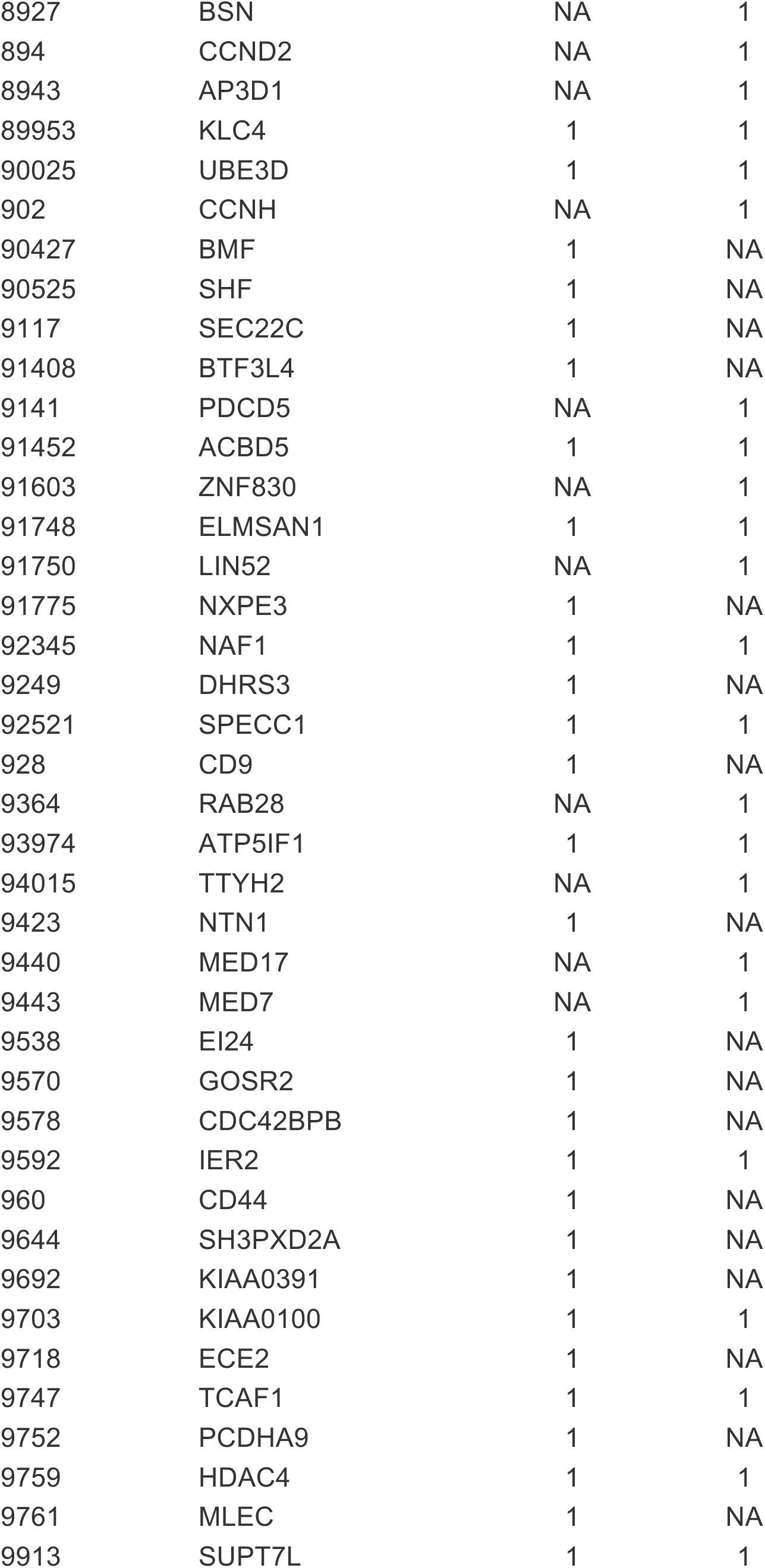
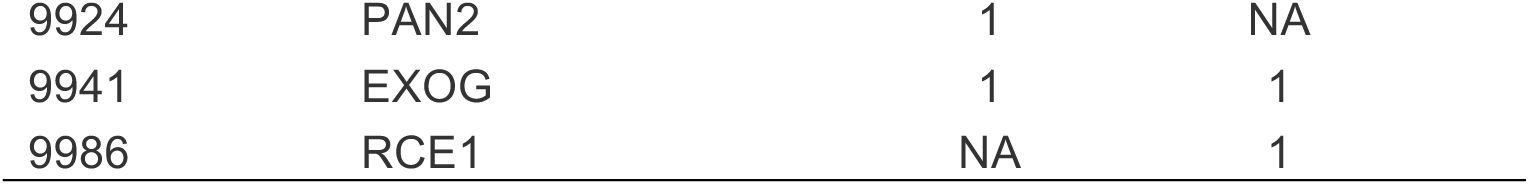
VMP significant gene/promoter hits identified in the MD affected versus MD unaffected contrast.

**Supplemental Table 4.**
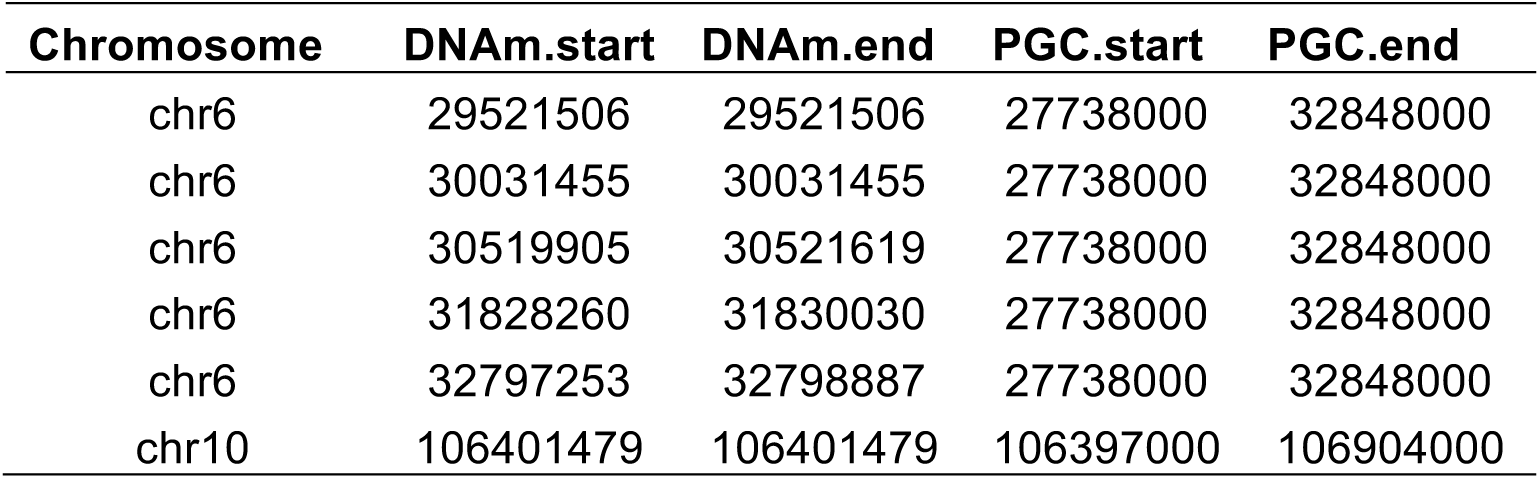
Overlap of MD related DMPs and DMRs with PGC GWAS loci.

| **Chromosome** | **DNA.m.start** | **DNA.m.end** | **PGC.start** | **PGC.end** |
| --- | --- | --- | --- | --- |
| chr6 | 29521506 | 29521506 | 27738000 | 32848000 |
| chr6 | 30031455 | 30031455 | 27738000 | 32848000 |
| chr6 | 30519905 | 30521619 | 27738000 | 32848000 |
| chr6 | 31828260 | 31830030 | 27738000 | 32848000 |
| chr6 | 32797253 | 32798887 | 27738000 | 32848000 |
| chr10 | 106401479 | 106401479 | 106397000 | 106904000 |

**Supplemental Table 5.** Overlap of MD related VMPs and VMRs with PGC GWAS loci.

| **Chromosome** | **DNA.m.start** | **DNA.m.end** | **PGC.start** | **PGC.end** |
| --- | --- | --- | --- | --- |
| chr2 | 58265673 | 58265673 | 57765000 | 58485000 |
| chr5 | 164658718 | 164658718 | 164440000 | 164789000 |
| chr6 | 28706472 | 28706472 | 27738000 | 32848000 |
| chr6 | 28890673 | 28890673 | 27738000 | 32848000 |
| chr6 | 31324972 | 31324972 | 27738000 | 32848000 |
| chr6 | 31707922 | 31707922 | 27738000 | 32848000 |
| chr6 | 31802397 | 31802397 | 27738000 | 32848000 |
| chr6 | 32049516 | 32049825 | 27738000 | 32848000 |
| chr6 | 32055738 | 32055738 | 27738000 | 32848000 |
| chr6 | 32789916 | 32789916 | 27738000 | 32848000 |
| chr6 | 32818212 | 32818212 | 27738000 | 32848000 |
| chr6 | 32847830 | 32847830 | 27738000 | 32848000 |

